# Analysis of Head and Neck Cancer scRNA-seq Data Identified PRDM6 Promotes Tumor Progression by Modulating Immune Gene Expression

**DOI:** 10.1101/2025.03.04.641548

**Authors:** Zhenyu Wu, Thurbu Tshering Lepcha, Dawei Zhou, Zhixian He, Guillaume N. Fiches, Youngmin Park, Jinshan He, Jianwen Chen, K.A.S.N Shanaka, Steve Oghumu, Weiqiang Zhao, Anjun Ma, Qin Ma, Jian Zhu, Netty G. Santoso

## Abstract

Head and neck squamous cell carcinoma (HNSCC) is a biologically aggressive and heterogeneous group of cancers with limited treatment options for patients who do not respond to standard therapies. While HPV-related HNSCCs tend to show better therapeutic outcomes, we still had limited understanding of the immune mechanisms underlying these cancers. Immune-responsive genes (IRGs) have emerged as critical factors in regulating both tumor progression and immune response. Recent advances in single-cell RNA sequencing (scRNA-seq) and the development of cell-type specific regulon inference tools, such as IRIS3, have provided new insights into the tumor immune microenvironment. In this study, we leveraged the IRIS3 platform to analyze scRNA-seq data from HNSCC patient samples, identifying novel transcription factor (TF)-IRG networks that contribute to tumor proliferation and immune escape. Specifically, we identified PRDM6, a histone methyltransferase, possesses the previously unknown role in promoting tumor cell proliferation by inducing IRG expression. We further demonstrated that HPV viral oncoproteins (E6/E7) oncoproteins up-regulate the PRDM6 expression, which associates PRDM6 with HPV-positive HNSCC.

## Introduction

Head and neck cancers (HNCs) represent the sixth most common cancer worldwide with the majority being head and neck squamous cell carcinoma (HNSCC). HNSCC typically arise from the mucosal linings of the upper aerodigestive tract (1, 2) and can be further classified according to its originating location, including the oral cavity, oropharynx, nasal cavity, paranasal sinuses, nasopharynx, larynx, and hypopharynx. Overall, HNSCC is highly heterogeneous and biologically aggressive, often associated with high rates of recurrence and mortality, especially at the advanced stages. Major risk factors of head and neck cancers include consumption of alcohol, exposure to nicotine, and infection with high-risk HPV (HPV16, 18) (*3*). Currently, the standard treatment of HNSCC includes surgery and/or chemo- and radiotherapy. Interestingly, HPV-associated HNSCC tends to have the better responses to treatments compared to HPV-negative ones (*4*), possibly due to its viral-related immunogenicity.

Despite of multiple anti-cancer treatment options, most of the locally advanced HNSCC cases still show poor responses with frequent recurrence. Immunotherapies have recently emerged as a promising alternative strategy to treat HNSCC considering that immune escape critically contributes to tumor initiation and progression (*5*). However, it is still at its early stage due to the lack of knowledge regarding the mechanism of immune regulation of HNSCC (*6*). Recently, it has been recognized that cancer cell-intrinsic genetic events profoundly modulate the tumor immune milieu and critically determine the outcome of immunotherapies (*7*). Immune-responsive genes (IRGs), thus, play a prominent role in not only controlling tumor initiation and progression but also participating in immune and inflammatory responses in tumor cells (*8, 9*). Moreover, several IRGs have been identified as tumor suppressors in various cancers, which directly impacts tumor growth (*10*).

Immunosurveillance plays an important role in controlling tumor initiation and progression. Malignant cells exploit various regulatory strategies to escape from immunosurveillance, including dysregulation of type I interferon (IFN) signaling (*11, 12*), immune checkpoint genes (*8*), as well as inflammatory responses (*13*) in tumor cells. Identification of TFs that govern IRG expression and depiction of related profound TF-IRG regulatory networks in tumor cells is critical to the understanding of tumor immunity, especially their own contribution directly from tumor cells. Through analysis of bulk RNA-seq datasets from The Cancer Genome Atlas (TCGA) consortium, differentially expressed IRGs and associated TF-IRG networks of HNSCC have been identified (*14, 15*). However, such bulk RNA-seq based analysis is unable to characterize cell type-specific IRG regulation.

The recent advance of single-cell RNA sequencing (scRNA-seq) technology enables the high-throughput analysis of gene expression at the resolution of individual cells. In our work, we developed a pipeline to perform the integrative analysis of multiple HNSCC scRNA-seq datasets and further the tumor cell-specific regulon inference analysis through the IRIS3 server (*16*). Results from our analysis provide a deeper understanding regarding the regulation of IRG expression in HNSCC malignant cells and the contribution of IRG dysregulation to tumor cell growth and immune response (*17*). In our studies, we utilized IRIS3 server to perform the integrative analysis of HNSCC scRNA-seq datasets, which led to the identification of tumor cell-specific regulons. Differentially expressed IRGs and novel transcription factor (TF)-IRG networks in tumor cells within the HNSCC microenvironment have been recognized. Here we report that PRDM6, a histone methyl-transferase (*18, 19*), which has never been implicated in HNSCC previously, regulate immune gene expression in HNSCC tumor cells, including interferon-stimulated genes (ISGs) ISG15 and IFITM1. Furthermore, we unraveled that HPV viral oncoproteins (E6/E7) induces PRDM6 expression, indicating PRDM6’s role in promoting HPV-positive HNSCC.

## Results

### IRIS3 analysis of HNSCC scRNA-seq data identified tumor cell-specific IRG-enriched regulons

We aimed to identify tumor cell-specific IRG-enriched regulons by using the publicly available HNSCC scRNA-seq datasets (**Table 1**). Our pipeline consisted of initial quality-control (QC) step through Seurat to filter out low quality cells. A set of established cell biomarkers was used to annotate tumor cells and non-tumor cells (primarily immune or stromal cells). Any available metadata containing cell annotation were used for confirmation. Datasets with a low percentage of tumor cells (less than 5%) were excluded from this study to avoid imbalances (**Fig. 1A**). In total, scRNA-seq data from fourteen HNSCC patients were included for analysis. Patient information, including viral infection status and clinical cancer stages, was summarized (**Table 1**). We performed inference of cell type-specific regulons using the IRIS3 tool. By leveraging public ChIP-seq data and bi-clustering methods, IRIS3 effectively identified the cell type-specific regulons with high accuracy and specificity. We were able to discover 639 TF-associated regulons that were specific for tumor cells in HNSCC tissues (**Table S1**). Since we primarily focused on immune-related regulons, we further performed the hypergeometric test to sort the regulons, in which IRGs were enriched. In total, we identified 88 tumor cell-specific TFs in the IRG-enriched regulons that were present in at least one patient (**Table S2**). Among these, 10 TFs in ≥ ten patients, 24 TFs in ≥ five patients, and 52 TFs in ≥ two patients (**Fig. 1B**). We highlighted those 24 TFs that occurred in more than 5 patients (**Fig. 1C**). Pathway analysis revealed that the regulons that associate with these 24 TFs are predominantly enriched in metabolism regulation and RNA processing (**Fig. S1A-C**), while IRG filtering indeed enriched the immune-related pathways for these regulons, including the response to viral infections and the NF-κB signaling (**Fig. S1A-C**). Among 24 TFs identified in more than five patients, several ones have been previously linked to HNSCC (*20–24*) (**Fig. 1D**), which validated the robustness of our IRIS3 analysis.

**Figure 1.**
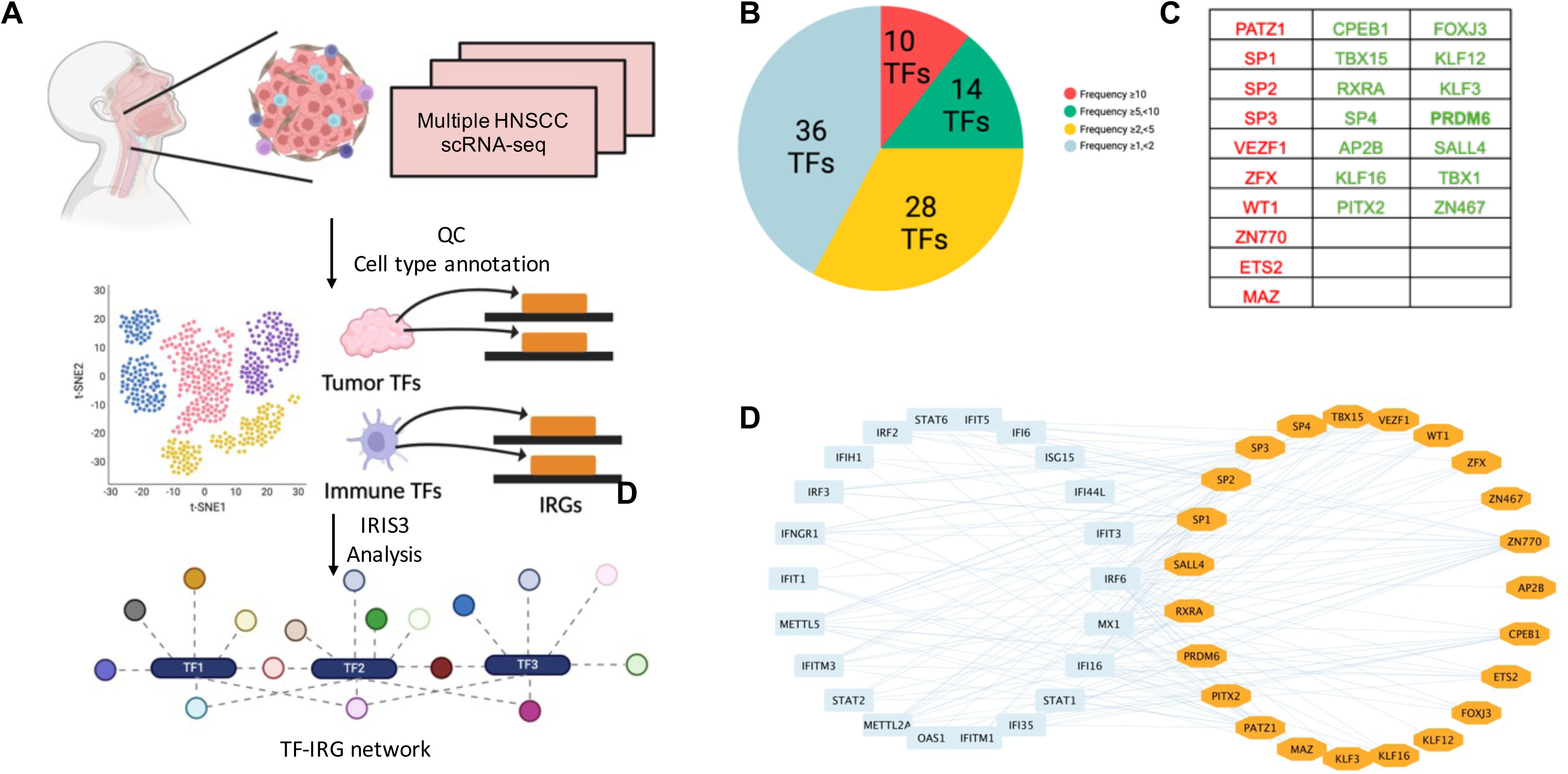
Identification of tumor cell-specific TF-IRG regulons through analysis of HNSCC scRNA-seq data. (A) The workflow utilized to identify tumor cell-specific TF-IRG regulons of HNSCC. HNSCC scRNA-seq datasets were collected and checked for quality. Clustering and annotation of single-cell data were performed by using Seurat. Cell-type specific regulons were identified by using the IRIS3 tool. (B) There are in total 88 TFs identified to regulate the expression of IRGs in HNSCC tumor cells. TFs were grouped according to their occurrence in multiple patients. (C) The 24 TFs identified in at least 5 patients were listed. (Red: TFs identified from ≥ 10 HNSCC patients; Green: TFs ≥ 5 HNSCC patients.) (D) Visualization of regulatory connections between 24 TFs and selected IRGs by using cytoscape (IRG: blue; TF: orange).

**Table 1.**
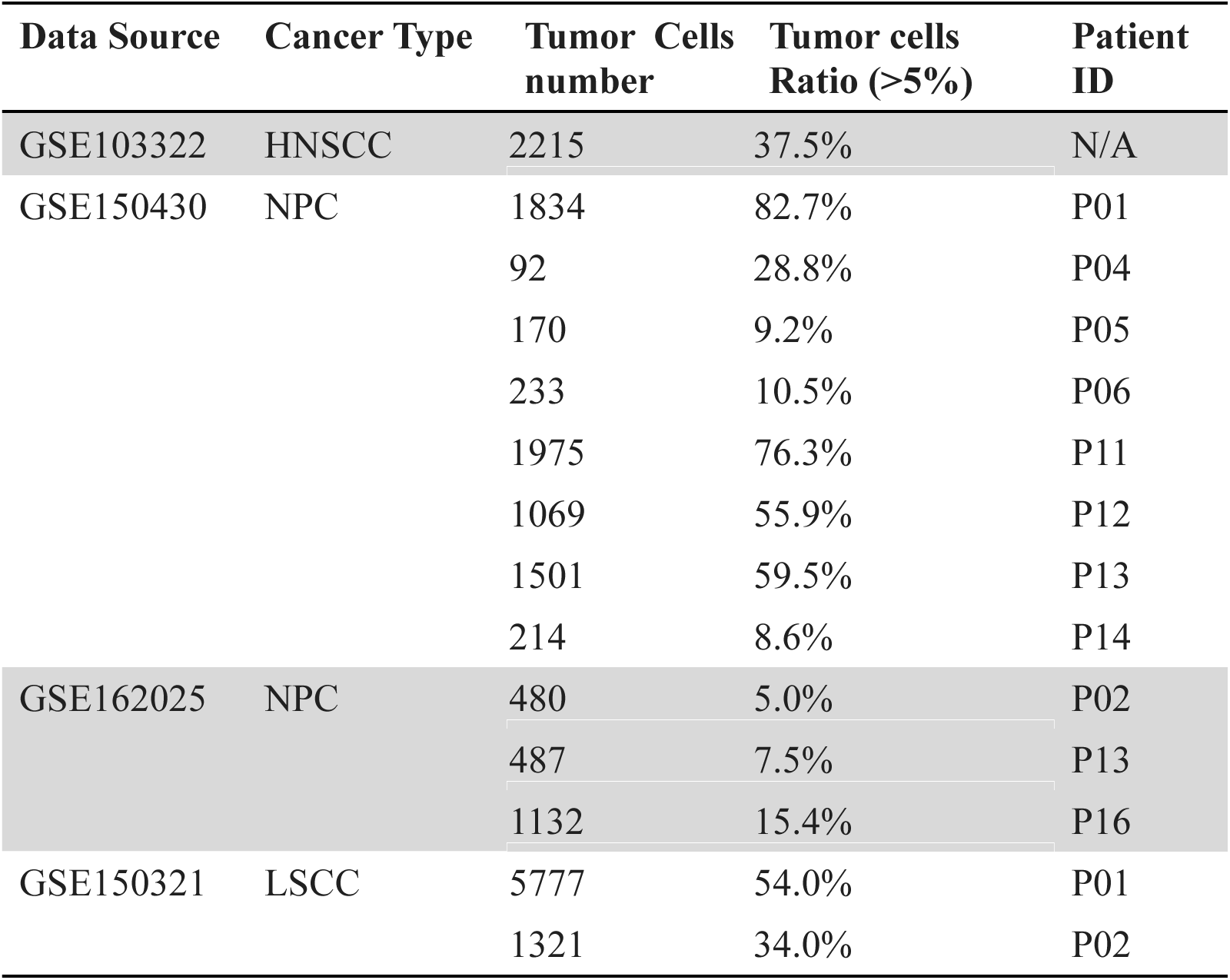
The list of HNSCC scRNA-seq datasets used in the study. The number and percentage of tumor cells were provided for each of 14 HNSCC patients. Certain patients were excluded from the analysis due to either insufficient number of tumor cells or low percentage.

### PRDM6 was recognized as a novel HNSCC-associated TF that regulates expression of IRGs in tumor cells

We focused on PRDM6 for further characterization since it has never been investigated in HNSCC. PRDM6 is a histone methyltransferase that regulates the methylation of H3K27 (*19*) and H4K20 (*25*). We identified 571 genes (**Table S3**) and 125 IRGs (**Fig. 2A**) in the PRDM6-associated regulon. To further investigate PRDM6’s role in regulating IRGs, we investigated the global chromatin associations of PRDM6 by re-analyzing the public ChIP-seq datasets for human PRDM6 (**Fig. S2A,B**). We observed that the peak distributions of PRDM6 are consistent across two ChIP-seq datasets, GSE76496 (*26*) and GSE106058 (*27*), and that PRDM6 occupies largely near the introns and distal intergenic regions but around 14% peaks indeed near the promoter regions. We further re-analyzed the CUT&RUN data of PRDM6 (GSE243557) (*19*), which identified that more PRDM6 peaks enrich near the promoter regions (**Fig. S2C**). This study also provided the H3K27me3 CUT&RUN data in the scenario of PRDM6 overexpression. Our re-analysis showed that the distribution of H3K27me3 peaks (CUT&RUN) is similar to that of PRDM6 peaks (ChIP-seq), indicating that PRDM6 functions as a H3K27me3 methyltransferase (**Fig. S2D**). We also employed DeSeq2 to identify differentially expressed genes (DEGs) either upregulated or downregulated due to PRDM6 overexpression in human neuroepithelial stem cells (GSE243554). Pathway analysis of these DEGs using ClusterProfiler revealed that they are enriched in the pathways related to the viral life cycle and various developmental processes (**Fig. S2E,F**). Interestingly, scRNA-seq data indicated that PRDM6 expression occurs almost exclusively in malignant cells while rarely in other cell types within the HNSCC tumors (**Fig. 2B**). We further determined the PRDM6 expression experimentally by using HNSCC cancer cell lines and tissue microarrays (TMAs). Our results showed that the PRDM6 expression is much higher in HNSCC cancer cell lines (CAL27, SCC9) in comparison to telomerase-immortalized normal human oral keratinocytes (OKF6/TERT-2, TIGK) at both protein and mRNA levels (**Fig. 2C,D**). We further confirmed that PRDM6 protein is present in TMAs of three HNSCC subjects (S1-S3) by using protein immunofluorescence staining assays (**Fig. 2E**). Overall, we could conclude that PRDM6 is a HNSCC-associated TF that preferentially expresses in HNSCC tumor cells.

**Figure 2.**
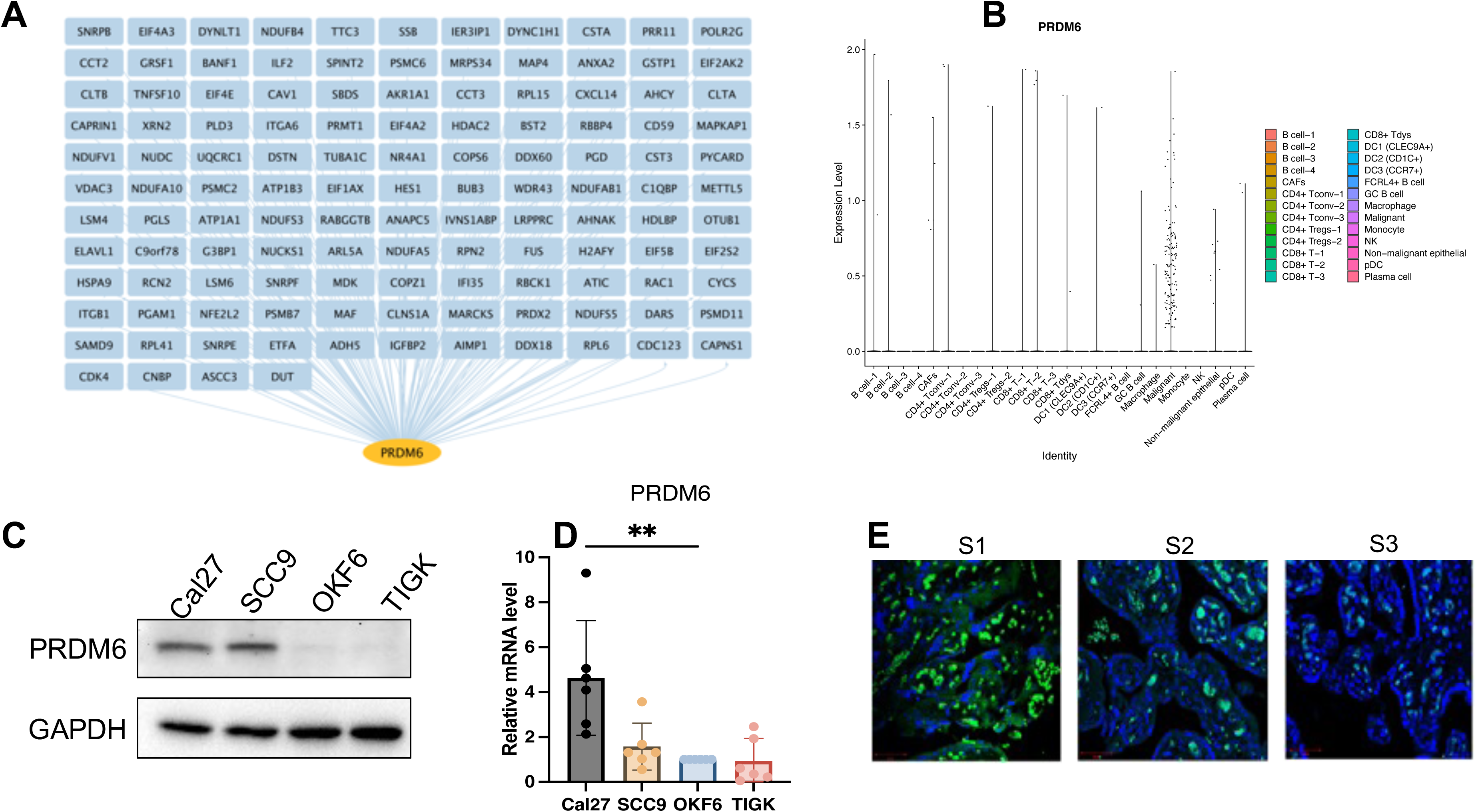
Characterization of PRDM6 as a HSNCC tumor cell-specific TF that regulates IRG expression. (A) PRDM6-IRG regulon identified by using the IRIS3 tool (IRG: blue; PRDM6: orange). (B) Expression level of PRDM6 across multiple cell types in HNSCC tumors from the public scRNA-seq dataset (GSE103322). Tumor and immune cells were annotated based on the expression of marker genes and internal labels. (C) Protein expression of PRDM6 in HNSCC cancer cell lines (CAL27, SCC9) as well as telomerase-immortalized normal human oral keratinocytes (OKF6/TERT-2, TIGK) was measured by protein immunoblotting analysis. GAPDH was used as the loading control. (D) The RNAs extracted from the above cells (C) were subjected to RT-qPCR analysis of PRDM6 transcript with normalization to GAPDH. (E) Expression of PRDM6 in tissue microarrays of three HNSCC subjects (S1-S3) was measured by protein immunofluorescence analysis with nuclei stained with Hoechst (PRDM6: green; nuclei: blue; scale bar: 50 μm). Results were calculated from two independent experiments and presented as mean ± SD. (*p < 0.05, **p < 0.01, ***p < 0.001, ****p < 0.0001, Student’s *t* test).

### PRDM6 promoted cell proliferation while suppressing immune gene expression of HNSCC tumor cells

As PRDM6 was shown to express in HNSCC tumor cells, we next determined whether PRDM6 contributes to their growth in vitro. We cloned the human PRDM6 cDNA in the pcDNA3.1 vector, which was transfected in CAL27 to generate cell clones. Our results revealed that the PRDM6-overexpressing CAL27 cell clones (1 and 2) consistently exhibit the higher cell viability and proliferation compared to vector only control by quantifying cellular ATP levels (**Fig. 3A**). Such effects were observed in these cells to be cultured continuously up to 5 days (**Fig. 3B**). We postulated that PRDM6 contributes to HNSCC tumor cell growth likely through dysregulation of immune gene expression based on our findings that PRDM6 is a HNSCC-associated TF that participates in the TF-IRG regulons identified from IRIS3 analysis. First, we observed that the expression of PRDM6 is moderately upregulated in CAL27 cells with IFN-α stimulation (**Fig. S3**). Second, we identified that depletion of PRDM6 by its siRNAs induces the expression of anti-tumor ISGs, ISG15 and IFITM1 (**Fig. 3C**). ISG15 was previously reported to inhibit the growth of tumor cells (*28, 29*) while inducing their cell death (*30*) by targeting the NF-kB and p53 signaling. The loss of IFITM1 was also shown to induce the cell cycle arrest (*31, 32*). Third, we also showed that PRDM6 overexpression consistently results in the decrease of ISG15 and IFITM1 expression in CAL27 cell clones (1 and 2) (**Fig. 3D**). Overall, these results indicated that PRDM6 might participate in the type I IFN signaling and control the expression of antitumor ISGs in HNSCC tumor cells, thus promoting their proliferation and growth.

**Figure 3.**
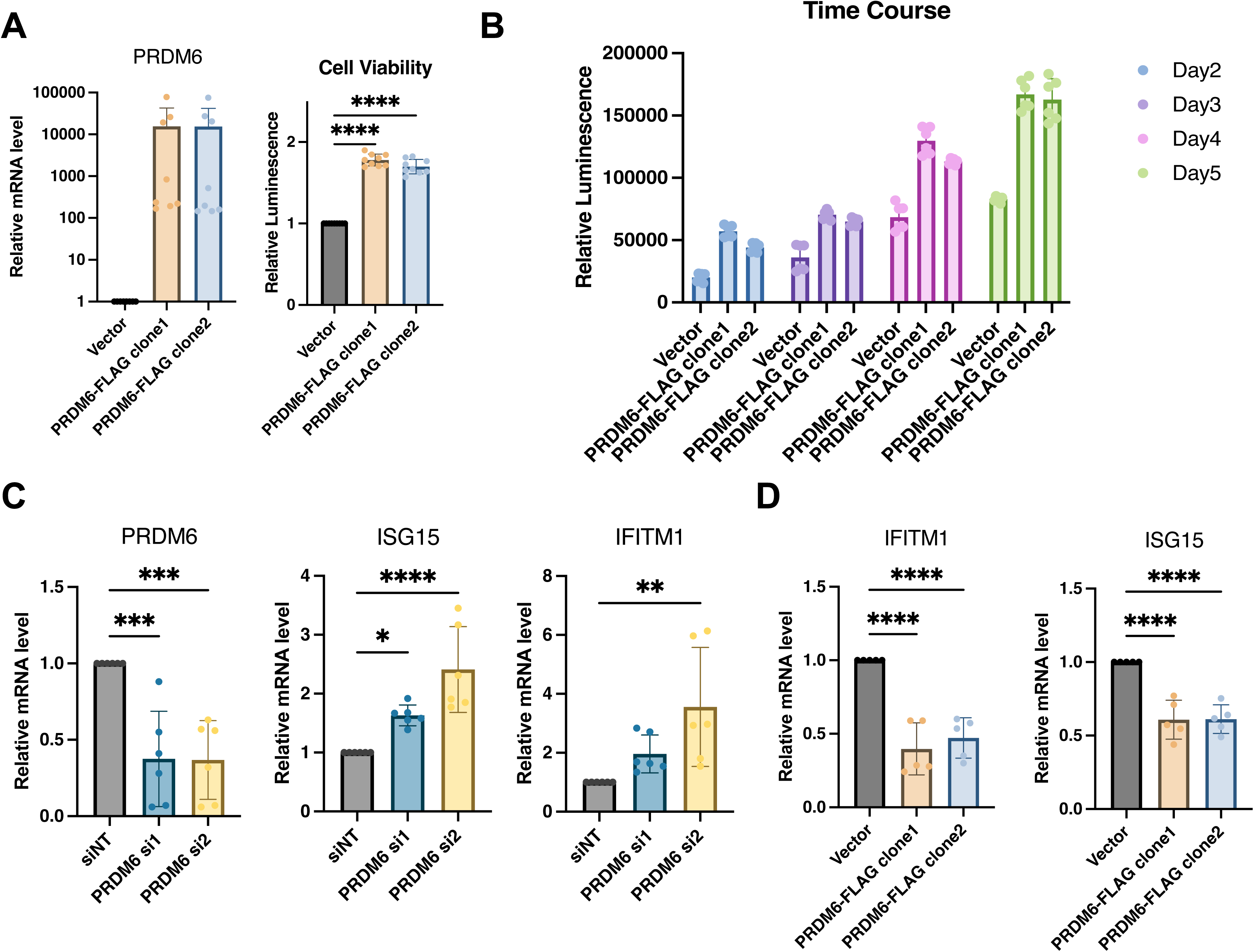
Functions of PRDM6 to promote growth of HNSCC tumor cells while suppress expression of anti-tumor ISGs. (A) The pcDNA vector expressing FLAG-PRDM6 was transfected in CAL27 cells to generate PRDM6-overexpressing cell clones (1 and 2). The empty vector (EV) was also transfected in CAL27 cells to generate a negative control. A portion of above cells was harvested for RNA extraction and RT-qPCR analysis of PRDM6 transcripts with normalization to GAPDH (left panel). The rest of cells was subjected to ATP-based cell viability analysis (right panel). (B) CAL27 cells transfected with the pcDNA vector expressing FLAG-PRDM6 (clones 1 and 2) or EV were cultured continuously. A portion of cells was harvested at the indicated timepoints and subjected to cell viability analysis. (C) siRNAs targeting PRDM6 (si1 and si2) or non-targeting control (siNT) were transfected in CAL27 cells. RNAs were extracted and subjected to RT-qPCR analysis of PRDM6 or ISG transcripts (ISG15, IFITM1) with normalization to GAPDH. (D) CAL27 cells transfected with the pcDNA vector expressing FLAG-PRDM6 (clones 1 and 2) or EV were subjected to RNA extractions and RT-qPCR analysis of ISG transcripts (ISG15, IFITM1) with normalization to GAPDH. Results were calculated from two independent experiments and shown as mean ± SD. (*p < 0.05, **p < 0.01, ***p < 0.001, ****p < 0.0001, Student’s *t* test).

### PRDM6 expression was upregulated by HPV-16 E6/E7 viral oncoproteins in HNSCC tumor cells

High-risk HPV (16 and 18) is a risk factor for HNSCC and specifically linked to oropharyngeal squamous cell carcinoma (OPSCC). HPV-encoded E6 and E7 (E6/E7) viral oncoproteins play a key role in driving the development and progression of HPV-associated cancers through various mechanisms (*33–35*). We next determined whether PRDM6 may participate in HPV E6/E7-mediated oral tumorigenesis. Our results showed that overexpression of HPV16 E6 and/or E7 oncoproteins indeed significantly increase PRDM6 expression in CAL27 cells by RT-qPCR analysis (**Fig. 4A**). We further confirmed this finding in the telomerase-immortalized normal human foreskin keratinocytes (N/Tert-1) that harbor the HPV16 genome (N/Tert-1+HPV16), showing that PRDM6 expression is higher in N/Tert-1+HPV16 compared to the HPV16-native N/Tert-1cells (**Fig. 4B**). Consistently, analysis of HNSCC bulk RNA-seq datasets from TCGA revealed that PRDM6 expression elevates in HPV-positive cases compared to HPV-negative ones (**Fig. 4C**), while expression of ISGs (ISG15, IFITM1) exhibits the opposite trend (**Fig. S4**). Overall, our findings implicated that HPV may target PRDM6 to suppress immune gene expression through induction of its expression by E6/E7 viral oncoproteins, thus promoting oral tumorigenesis (**Fig. 4D**).

**Figure 4.**
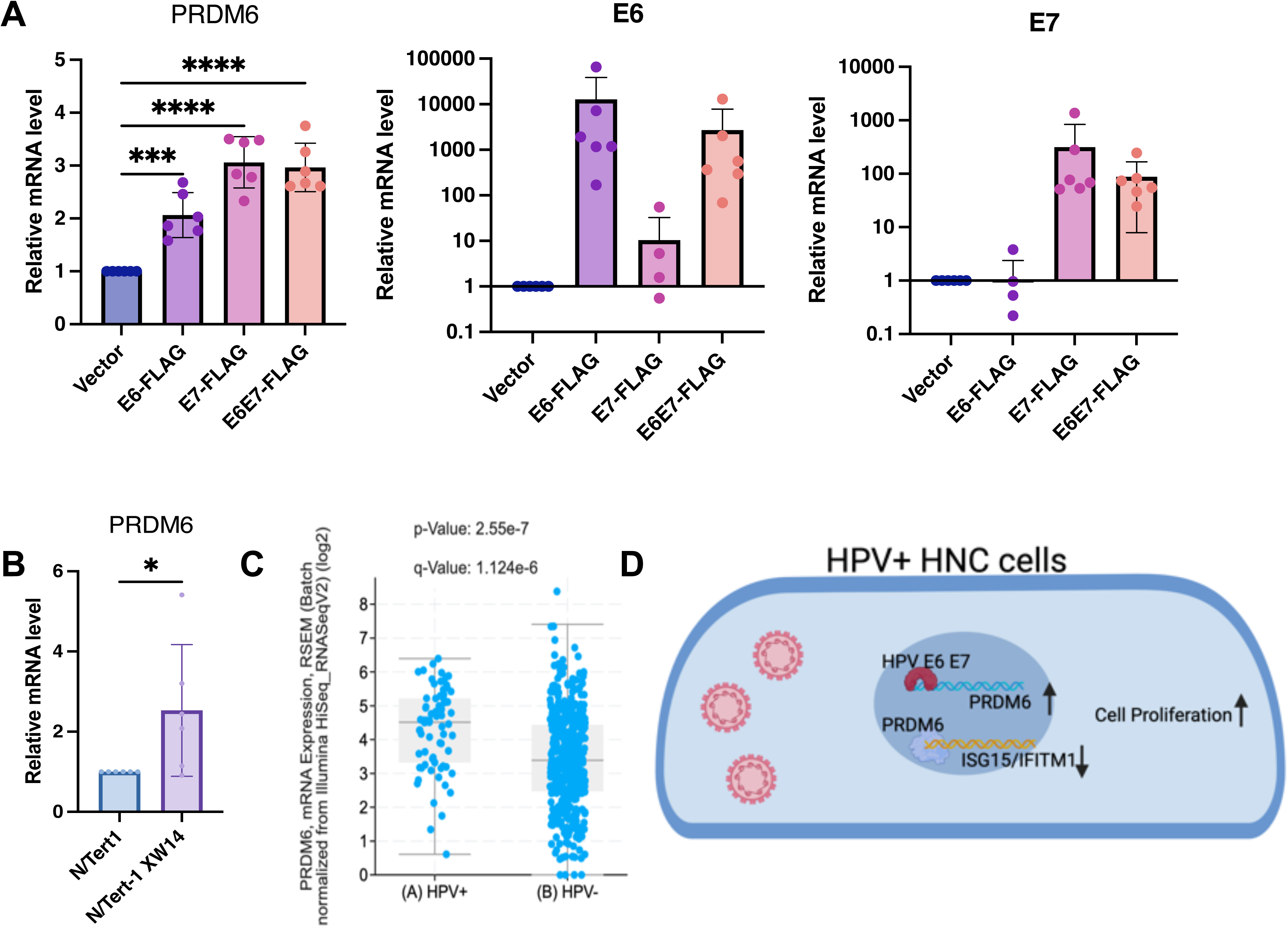
Induction of PRDM6 expression by HPV-16 E6/E7 viral oncoproteins in HNSCC tumor cells. (A) The pLXSN vector expressing FLAG-E6 (left panel), FLAG-E7 (middle panel), FLAG-E6/E7 (right panel), or the or empty vector (EV) was transfected in Cal27 cells. RNAs were extracted and subjected to RT-qPCR analysis of PRDM6 or E6/E7 transcripts with normalization to GAPDH. (B) The telomerase-immortalized normal human foreskin keratinocytes with or without HPV-16 genome, N/Tert-1+HPV16 or N/Tert-1 cells, were subjected to RNA extractions and RT-qPCR analysis of PRDM6 transcripts with normalization to GAPDH. (C) HNSCC bulk RNA-seq datasets from TCGA were re-analyzed. Expression levels of PRDM6 in HPV-positive and negative (HPV+/-) HNSCC patients were compared. (D) A working model described that HPV may induce PRDM6 expression by E6/E7 viral oncoproteins to suppress expression of anti-tumor ISGs and thus promote development of HNSCC. Results were calculated from three independent experiments and shown as mean ± SD. (*p < 0.05, **p < 0.01, ***p < 0.001, ****p < 0.0001, Student’s t test).

## Discussion

In this study, we integrated multiple datasets of HNSCC scRNA-seq with the aim to identify tumor cell-specific transcriptional/epigenetic regulators that control immune gene expression and tumor cell intrinsic immunity. Such efforts led to the identification of 639 tumor-specific TF regulons, including 88 IRG-enriched ones. Among them, the histone methyltransferase PRDM6 emerged as a top, previously un-investigated TF that may regulate the immune response in HNSCC tumor cells. We demonstrated that PRDM6 expression occurs almost exclusively in tumor cells of the clinic HNSCC tissues and that PRDM6 promotes proliferation and growth of HNSCC tumor cells *in vitro* while suppressing expression of anti-tumor ISGs (ISG15, IFITM1) by both loss- and gain-of-function approaches. We further identified that HPV16 E6/E7 oncoproteins upregulate PRDM6 expression in HNSCC tumor cells, which may link PRDM6 to HPV-induced oral oncogenesis.

TF gene regulatory networks as well as differentially expressed IRGs have been identified through analysis of HNSCC bulk RNA-seq datasets from TCGA (*36–38*). However, inherent limitations of bulk RNA-seq forbid the high-resolution characterizations of cellular heterogeneity within the tumor microenvironment, as gene expression levels across diverse types of cells in the tumor tissues are quantified with the setting that all types of cells are mixed together. Additionally, the regulon predictions primarily rely on the previous knowledge of benchmarked connections between TFs and their gene targets (*39, 40*). On the contrary, the scRNA-seq based transcriptomic profiling overcomes these limitations of bulk RNA-seq and enables the precise measurements of gene expressions that occur exclusively in malignant cells *vs* other types of cells in the tumor tissues, which permits us to annotate the HNSCC tumor cell-specific TF-IRG regulatory networks with the higher accuracy. IRIS3, powered by its unique bi-clustering algorithm and *de novo* motif prediction, has demonstrated its capability to identify the cell type-specific regulons with the satisfying reproducibility and robustness (*16*), which enabled our successful identifications of 88 IRG-enriched TFs in the HNSCC tumor cells (**Fig. 1**).

We further characterized the histone methyltransferase PRDM6 as a potential TF that governs HNSCC tumor cell intrinsic immunity. Beyond the mere data mining, we experimentally verified that PRDM6 expresses in both HNSCC cancer cell lines and HNSCC clinic TMAs from multiple subjects (**Fig. 2**). We also demonstrated PRMD6’s functions to promote proliferation and growth of HNSCC tumor cells, likely due to its activities to suppress expression of anti-tumor ISGs (**Fig. 3**). These results are overall consistent with the early findings that PRDM family of proteins possess the oncogenic potency, and regulate the tumor cell proliferation and differentiation (*18, 19, 25, 41, 42*). However, the exact underlying mechanisms for PRDM6 to drive development and progression of HNSCC still need further investigations in future. Nevertheless, it has been showcased that PRDM6 indeed targets the histone marks including H3K27 in medulloblastoma (*19*). It is thus plausible that PRDM6-induced H3K27 trimethylation (H3K27me3) likely leads to the suppression of immune gene expression in tumor cells.

We also reported that PRDM6 suppresses immune gene expression in HNSCC tumor cells, which is consistent with the previous studies on other members of PRDM proteins. In particular, PRDM1 is known to play a key role in immunosuppression in the tumor microenvironment. It has been shown that genetic ablation of PRDM1 is effective to stimulate antitumor T cell responses (*43*). On the contrary, the immunosuppressive functions of PRDM6 need to be further characterized. We did identify that PRDM6 expression is inducible in HNSCC tumor cells upon type I interferon stimulation, indicating that PRDM6 may be involved in tumor cell intrinsic immune signaling. There are also other clues. For example, it has been noted that PRDM6 protein associates with the histone methyltransferase G9a that is a critical regulator of inflammatory responses as well as T cell functions (*44*). Overall, we provided the evidence that PRDM6 suppresses expression of antitumor ISGs, which may contribute to oral tumorigenesis.

HPV highly associates with HNSCC, especially OPSCC, as nearly 80% of oropharyngeal cancers in the United States are infected with the high-risk HPV (16 and 18) (3). HPV E6/E7 viral oncoproteins are critical to drive malignant transformation of normal oral epithelial cells by targeting key cellular pathways with multiple strategies, such as degradation of p53 protein (*45–47*), activation of human telomerase reverse transcriptase (hTERT) (*48, 49*), and disruption of retinoblastoma (Rb) protein (*50–52*). Our results suggested a new link that HPV E6/E7 may manipulate PRDM6 expression to antagonize the antitumor and antiviral immune responses as a previously unappreciated viral mechanism to promote oral tumorigenesis (**Fig. 4**). These findings would improve the fundamental understanding of HPV-associated HNSCC. However, it remains to be delineated how E6/E7 transcriptionally or epigenetically induces PRDM6 expression.

To summarize, our unbiased analysis of integrated HNSCC scRNA-seq datasets led to the identification of PRDM6 as a novel TF that governs the type I IFN signaling and immune gene expression and thus promotes proliferation and growth of tumor cells, while PRDM6 itself is also under control of HPV. These results would shed light in targeting PRDM6 for developing novel therapies to treat HNSCC. The potential immunosuppressive functions of PRDM6 would likely need to be interrupted or counteracted to boost antitumor immune responses, thus improving the efficacy of immunotherapies against HNSCC.

## Material and Methods

### Cells

CAL27 and SCC9 cells were cultured in DMEM supplemented with 10% FBS. OKF6/TERT-2 (*53*) and TIGK cells (*53*) were both cultured in keratinocyte serum-free medium (K-sfm) with supplements (Thermo Scientific, CAT#17005042). Cell proliferation and growth was measured by using the ATP-based CellTiter-Glo Luminescent Cell Viability Assay (Promega, Cat. # G7572) following the manufacturer’s instructions and analyzed by the Cytation 5 multimode reader (luminescent mode).

### Quantitative PCR (qPCR)

Total RNAs were extracted using the NucleoSpin RNA extraction kit (Macherey-Nagel, Cat. # 740955.250) following the protocols provided by the manufacturer. RNA samples were reverse transcribed to cDNAs using iScript (BioRad, Cat. # 1708891). Real-time qPCR was performed on a CFX96 instrument (BioRad), by mixing 5 ul (2x) SYBR Green Supermix (BioRad, Cat. # 1725214), 0.5 ul primer mix, and 4.5 ul cDNA template. Data were analyzed by using the ΔΔCt method with GAPDH as an internal control. All qPCR primers were listed in **Table S4**.

### Protein immunoblotting

Protein immunoblotting was performed as previously described (*54*). The following antibodies were used: anti-PRDM6 (Invitrogen, Cat. # PA5-43659), anti-GAPDH (Santa Cruz Biotechnology, Cat. # sc-47724), anti-mouse-HRP (Cell Signaling technology, Cat. # 7076), anti-rabbit-HRP (Cell Signaling technology, Cat. # 7074). The membrane was washed and blocked with 5% BSA, followed by the incubation with the primary antibody (1:1000 dilution) in 10 ml of the antibody dilution buffer for overnight at 4 °C with shaking, and with the secondary anti-mouse or anti-rabbit antibody.

### Transfection and electroporation

Turbofect reagents (Thermo Scientific, Cat. # R0531) were used for plasmid transfection following the manufacturer’s recommendation as described previously (55). The pcDNA3.1 plasmid expressing the PRDM6 cDNA with FLAG tag at 3’ end was purchased from GenScript. The pLXSN plasmid expressing FLAG-tagged E6 (Cat. # 52395), E7 (Cat. # 52396), E6/E7 (Cat. # 52394) were acquired from Addgene. siRNAs targeting PRDM6, siPRDM6-1 (Cat. # s41097) and siPRDM6-2 (Cat. # s41098), were purchased from Invitrogen. For siRNA transfection in CAL27 and SCC9 cells, reverse transfection was performed using the Lipofectamine RNAiMAX (Invitrogen, Cat. # 13778100) as previously described (*9*).

### Protein immunofluorescence

Paraffin-embedded HNSCC tissue slides were baked at 65°C for 30 mins, followed by deparaffinization in xylene (10 mins × 2), 100% ethanol (10 mins × 2), 95% ethanol (5 mins), 70% ethanol (1 min), 50% ethanol (1 min), and finally rehydrated in double-distilled water (5 minus). Antigen unmasking was performed using 2100 Retriever (Electron Microscopy Sciences, Cat. # 62700-10) with the Antigen Retrieval Buffer (100X Tris-EDTA Buffer, pH9.0; Abcam, Cat. # ab93684) for 2 hrs to complete the cycle and then cool down. To block non-specific binding, slides were incubated with 10% normal goat serum (NGS) in PBST (0.1% Tween-20 in PBS) for 2 hrs at room temperature (RT). Incubation with the primary antibody was carried out overnight at 4°C using rabbit anti-PRDM6 antibody with 5% NGS in PBST. Sections were then washed with PBST and incubated with Alexa Fluor 488-conjugated, anti-rabbit secondary antibody with 5% NGS for 1 hr at RT. Nuclear counterstaining was performed using Hoechst 33342 (1:5000 in D-PBS) for 15 mins at RT. Coverslips were mounted on slides using ProLong Glass Antifade Mountant (Invitrogen, Cat. # P36982) and dried in the dark overnight. Confocal images were acquired using the ZEISS LSM 700 Upright laser scanning confocal microscope and processed with ZEN imaging software (ZEISS).

### Data analysis

Fastq files of the public bulk RNA-seq dataset (GSE243554) were obtained from GEO database and reanalyzed. Quality of raw reads was assessed by fastp (*56*). Reads were trimmed with adapters and aligned to the human genome (GRCh38.p14) with HISAT2 aligner. Uniquely aligned reads were submitted as inputs for mapping to gencode.v39 annotation. For identification of DEGs, DESeq2 was run with raw read counts obtained from FeatureCounts. Adjusted p value =< 0.05 and fold change >= 2 were used to filter for DEGs. For re-analysis of HNSCC scRNA-seq datasets, expression matrix was downloaded from GEO with the accession number GSE103322 (*57*), GSE150430 (*58*), GSE150321 (*59*), GSE162025 (*60*), and GSE150825 (*61*). Low-quality cells or empty droplets were filtered out based on the number of unique genes, the total number of molecules, and the percentage of mitochondrial reads detected in each cell. Normalization and scaling of cell clustering and visualization were performed using SeuratV4.0 (*62*). Cell identity was assigned based on the metadata and marker genes. Raw data of the public ChIP-seq datasets (GSE76496, GSE106058) were acquired from GEO and processed with ENCODE ChIP-seq-pipeline2. PRDM6 and H3K27me3 CUT&RUN peaks from GSE243557 were visualized using IGV. The code used for above analyses was deposited at github: https://github.com/ZhenyuWu-OSU

### Statistics

Statistical analyses of experiment results were performed by using the unpaired, two-tailed Student’s *t* test or the one-way analysis of variance (ANOVA) in Graphpad PRISM 9.0 package. Variables were compared among all tested groups, and the P values below 0.05 were considered as statistically significant.

## Acknowledgment

We would like to thank Dr. James G. Rheinwald for providing the OKF6/TERT-2 cells, and Dr. Jens Kreth (Oregon Health & Science University) for the TIGK cells. We appreciate the supports from Drs. Yuzhou Chang and Cankun Wang for performing the IRIS3 analysis. We also appreciate the supports from Dr. Meng Wang (Research Institute at Nationwide Children’s Hospital) to establish the analytic pipelines of ChIP-seq data. This study was supported by NIH research grants R01CA260690 to N.S.; R01MH134402, R01DA059538, R56AI181631 to J.Z.

## Supplemental Materials

**Figure S1.**
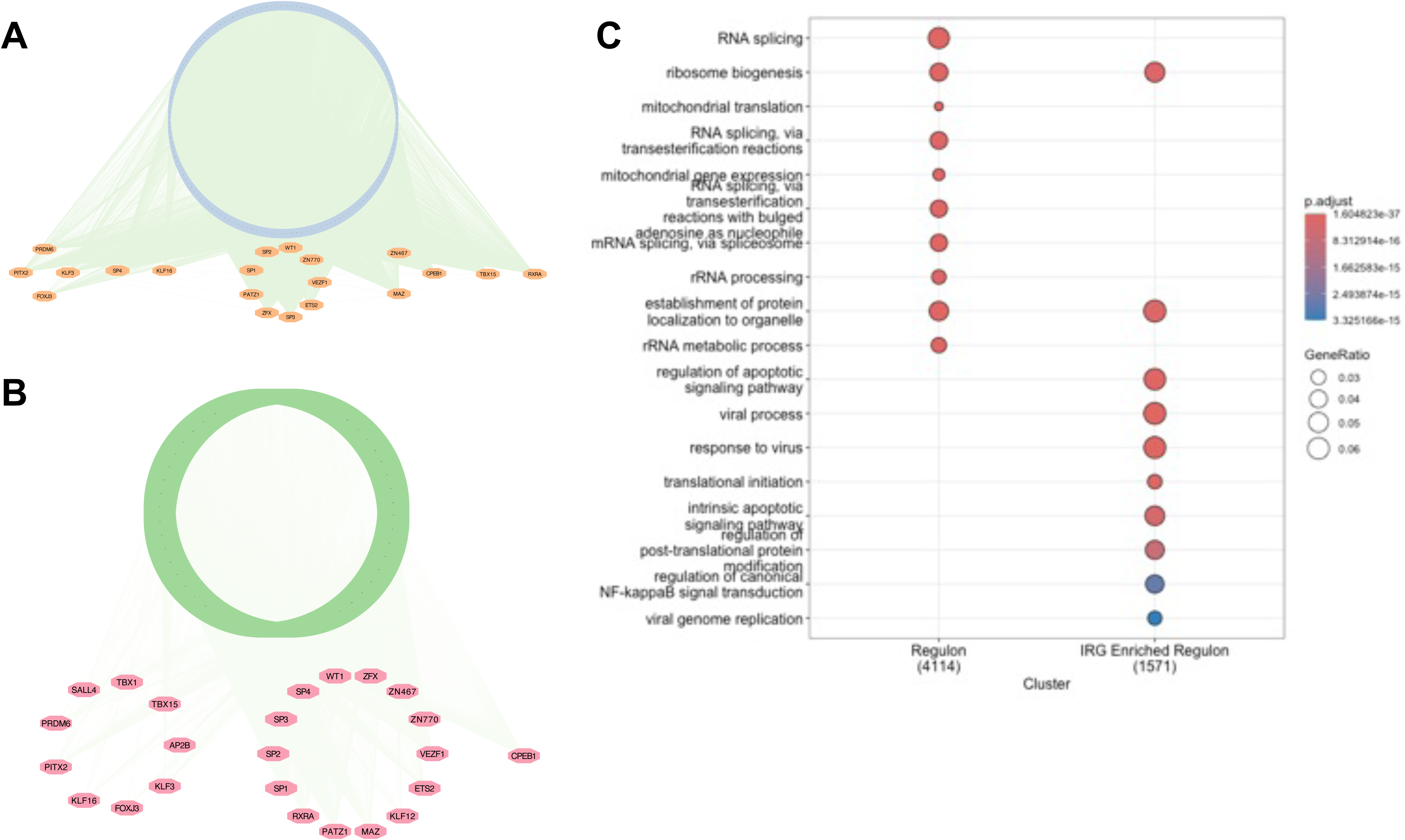
(A) Visualization of the TF-associated regulons for those TFs identified from at least 5 HNSCC patients. In total, 4,560 genes (blue) were predicted to under regulation of 24 TFs (orange), resulting in 36,696 predicted regulatory interactions between TFs and gene targets. TFs were clustered according to their eccentricity using Cytoscape. (B) Visualization of the TF-associated, IRG-enriched regulons for 24 TFs. The 4,560 genes were further filtered with the IRGs. In total, 1,636 IRGs (green) were predicted to under regulation of 24 TFs (red), resulting in 12,029 predicted regulatory interactions between TFs and IRGs. TFs were clustered according to their eccentricity using Cytoscape. (C) Pathway analysis was conducted for genes regulated by 24 TFs using ClusterProfiler, showing the enrichment of RNA processing pathway for general gene targets or viral response, transcription, and translation regulation pathways for IRGs, respectively.

**Figure S2.**
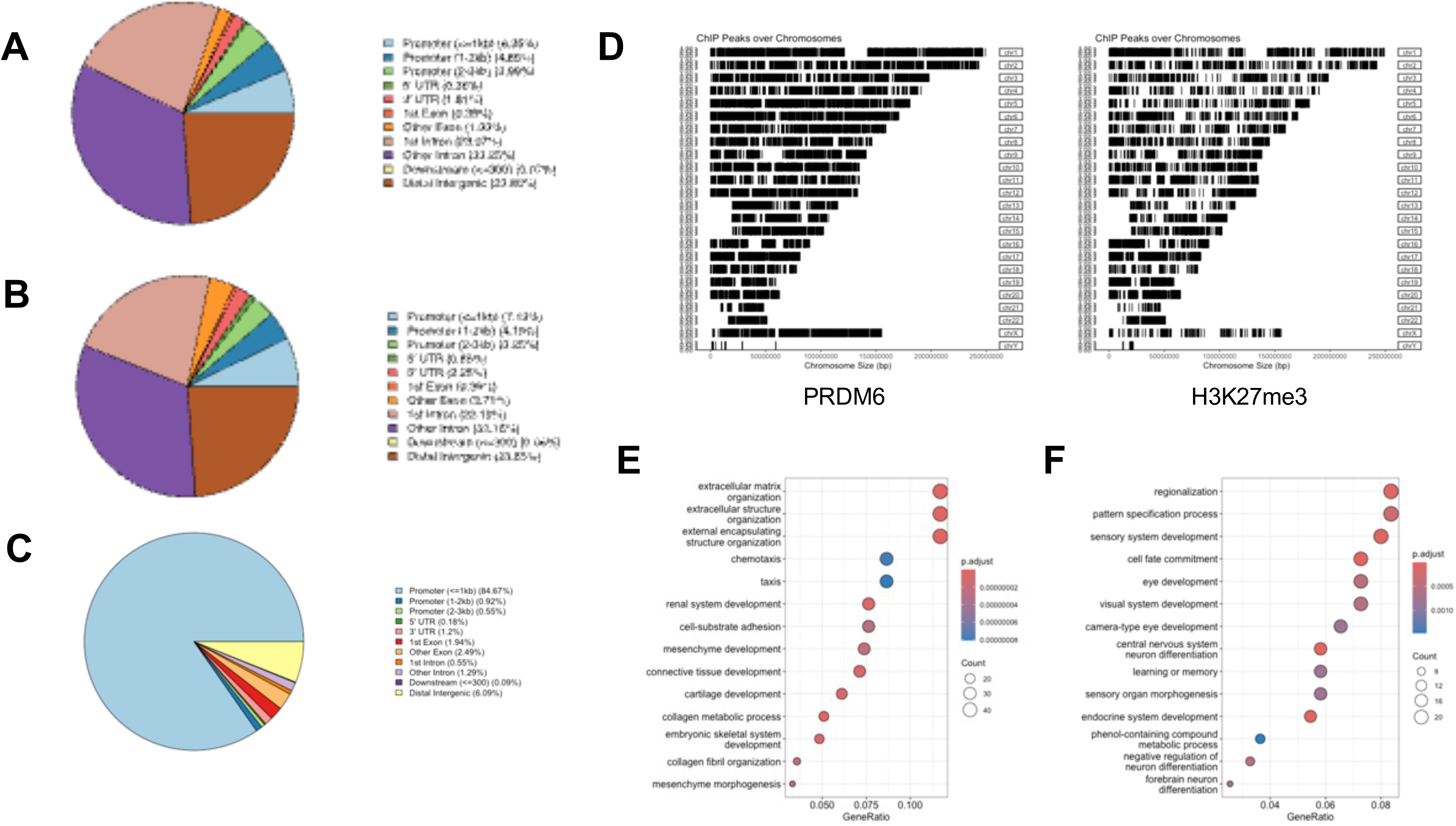
(A-C) Public PRDM6 ChIP-seq datasets GSE106058 (A) and GSE76496 (B), as well as PRDM6 CUT&RUN dataset GSE243557 (C) were collected for re-analysis. The distribution of PRDM6 binding peaks across human genomes was visualized using Chipseeker. (C) Re-analysis of public PRDM6 ChIP-seq data (GSE76496) and H3K27me3 CUT&RUN data (GSE243557) demonstrated the similar distribution of PRDM6 and H3K27me3 bindings peaks. (E, F) Pathway analysis of the upregulated (F) and downregulated (G) gene expression in human neuroepithelial stem (NES) cells with PRDM6 overexpression (GSE243554).

**Figure S3.**
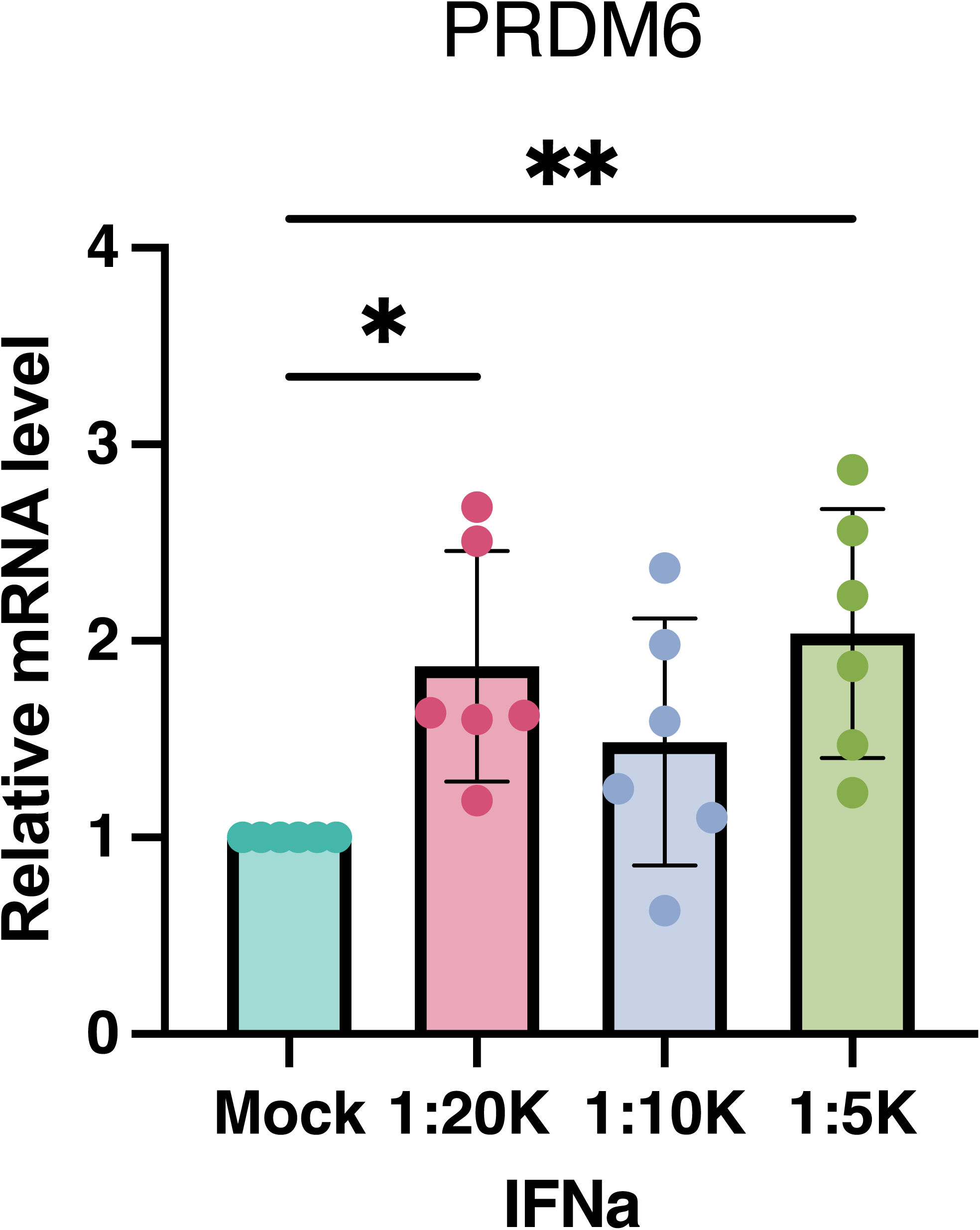
CAL27 cells were treated with IFN-α at the indicated doses. RNAs were extracted and subjected to RT-qPCR analysis of PRDM6 transcripts with normalization to GAPDH.

**Figure S4.**
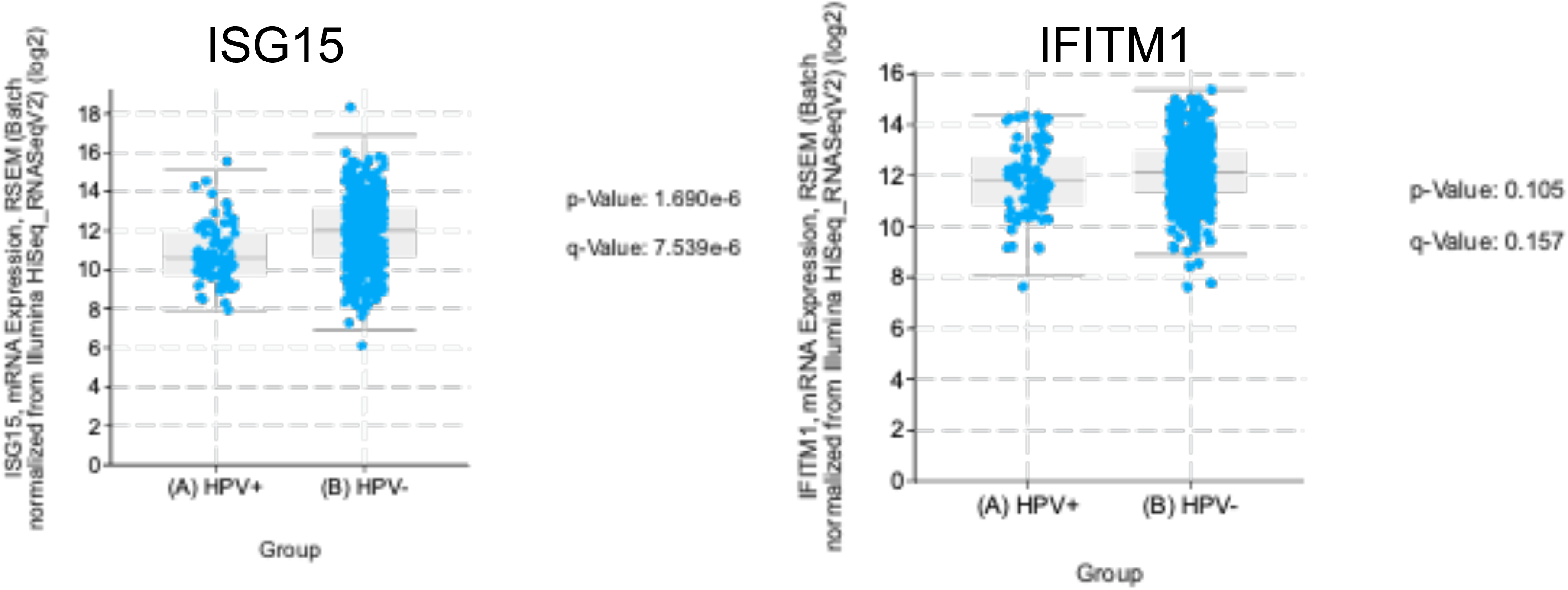
HNSCC bulk RNA-seq datasets from TCGA were re-analyzed. Expression levels of ISG15 and IFITM1 in HPV+/− HNSCC patients were compared.

**Table S1.**
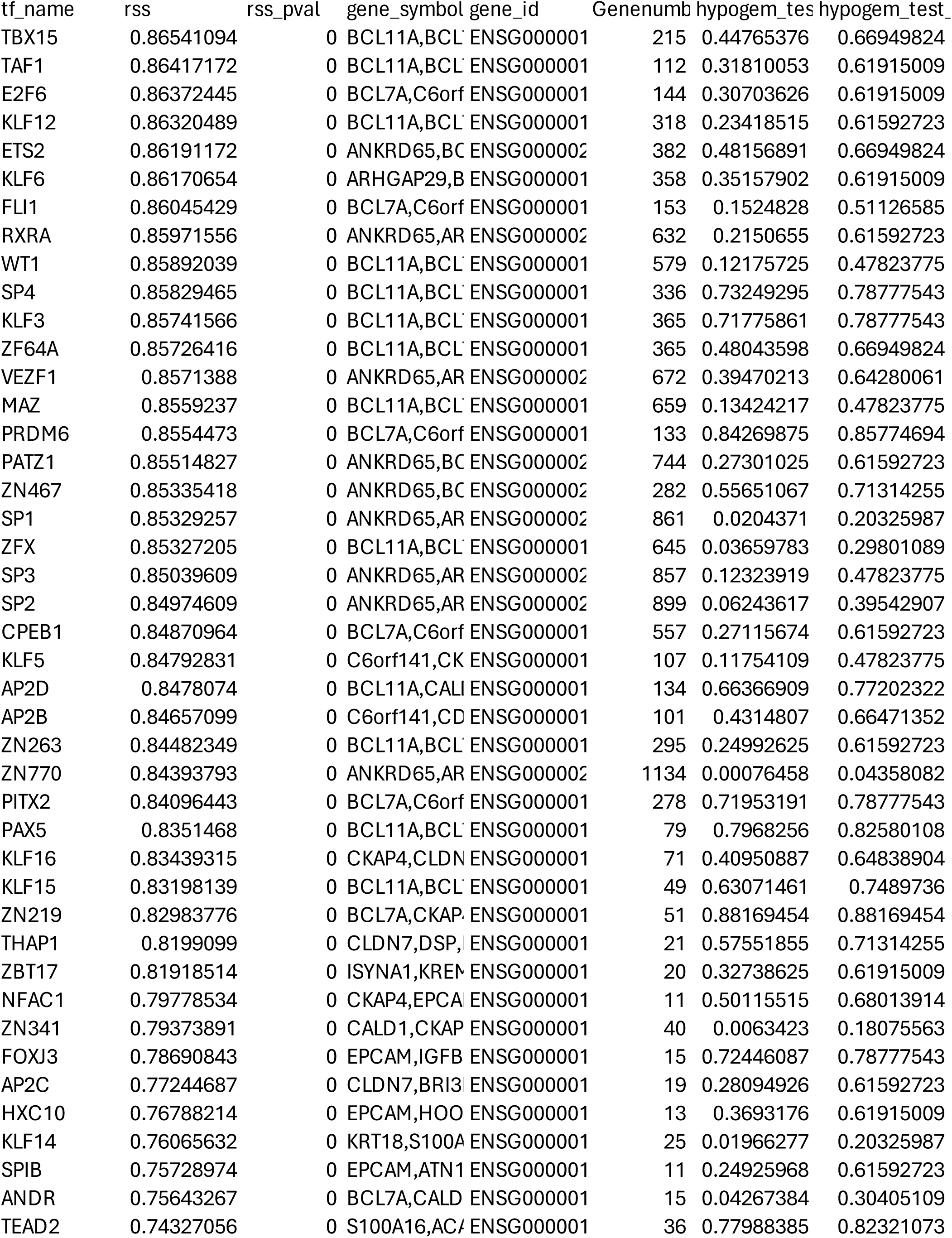
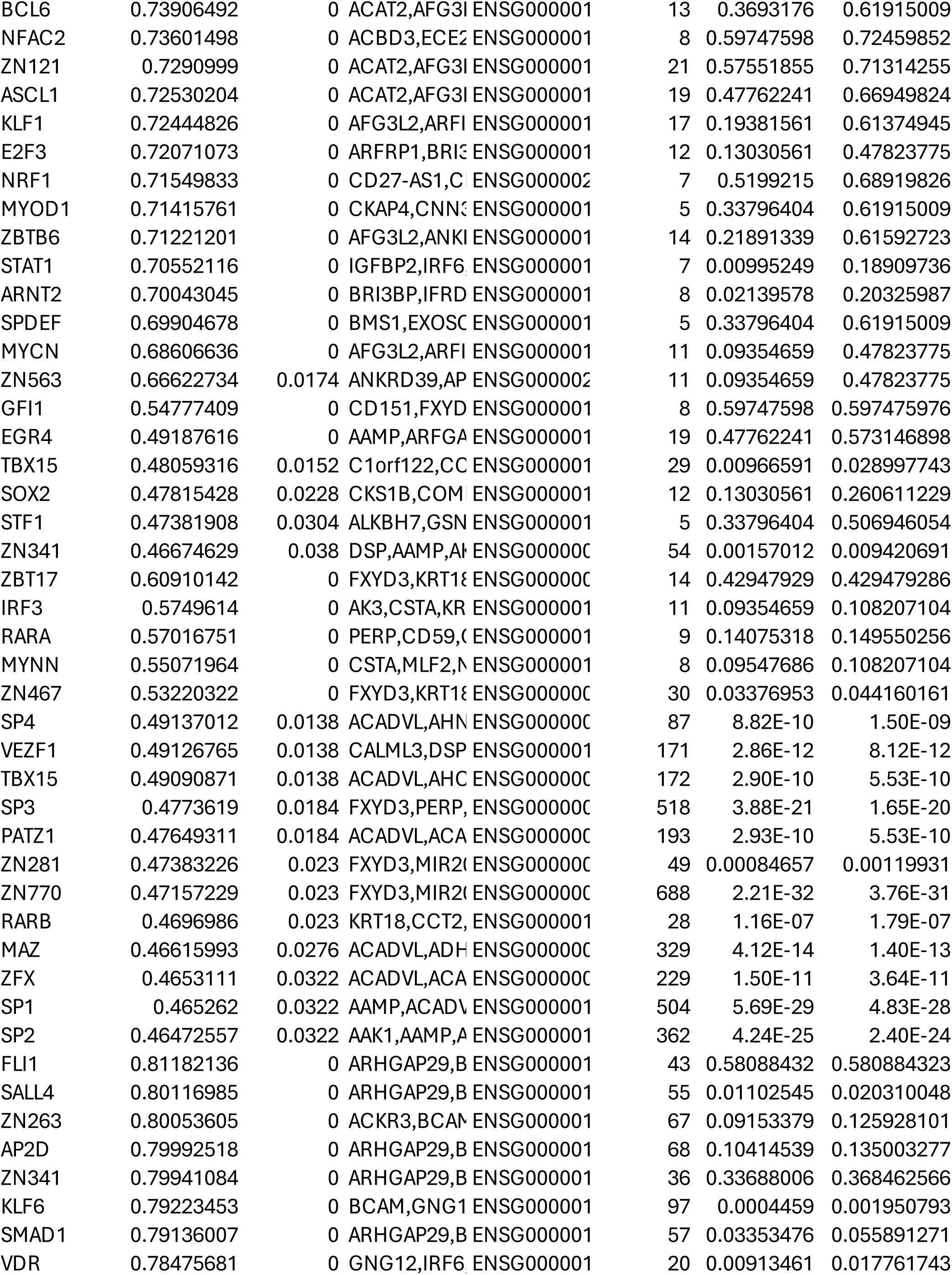
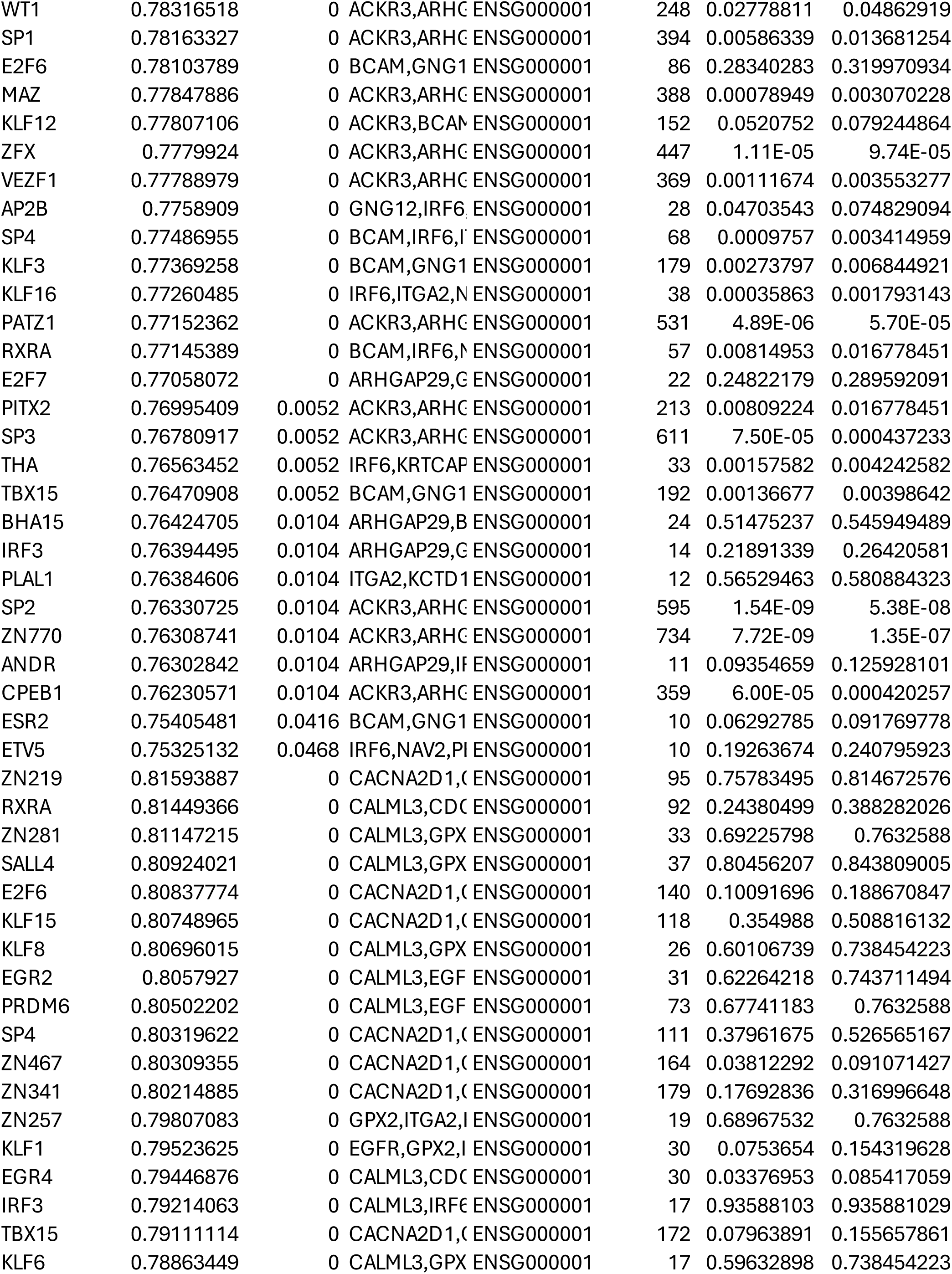
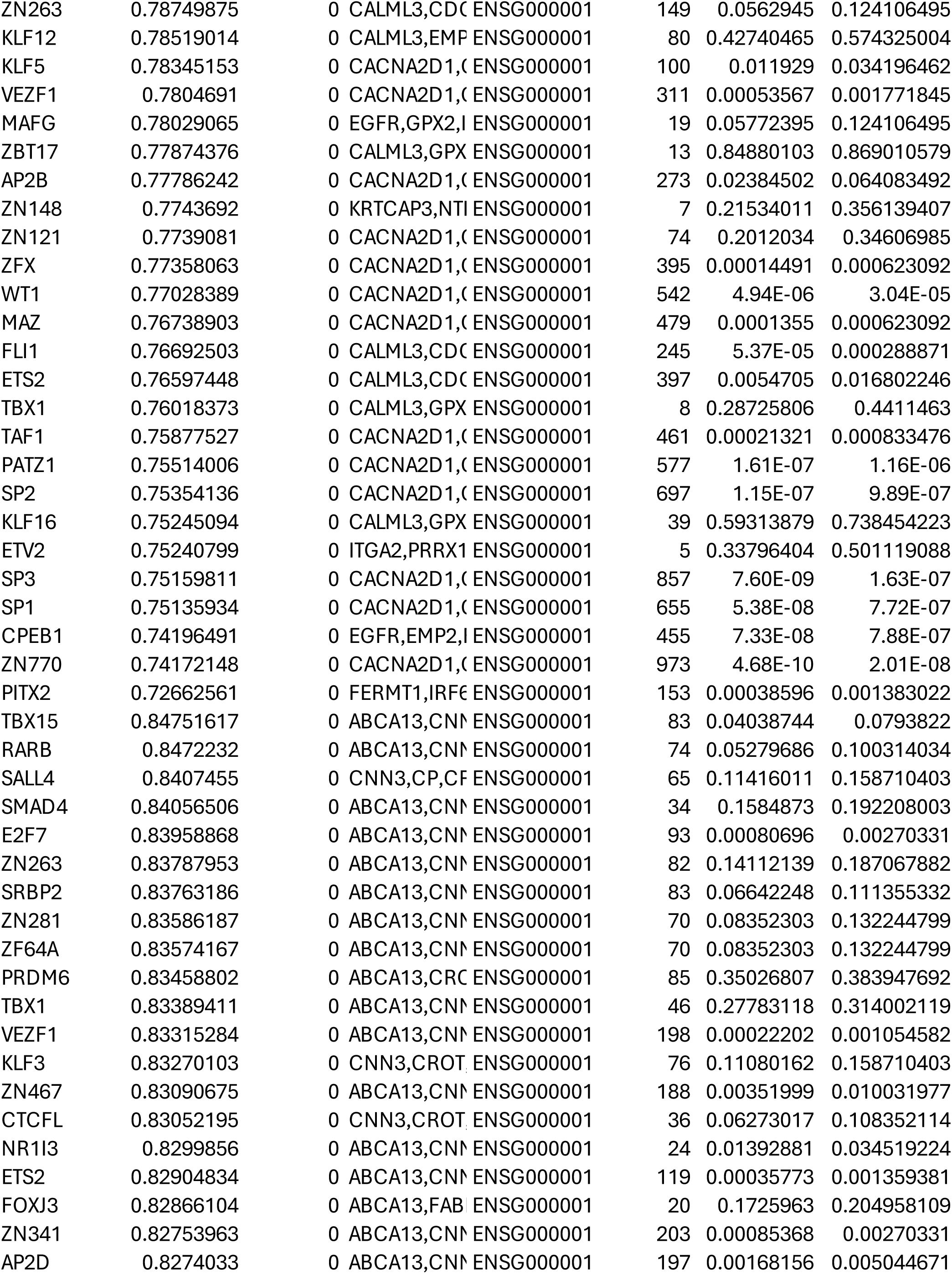
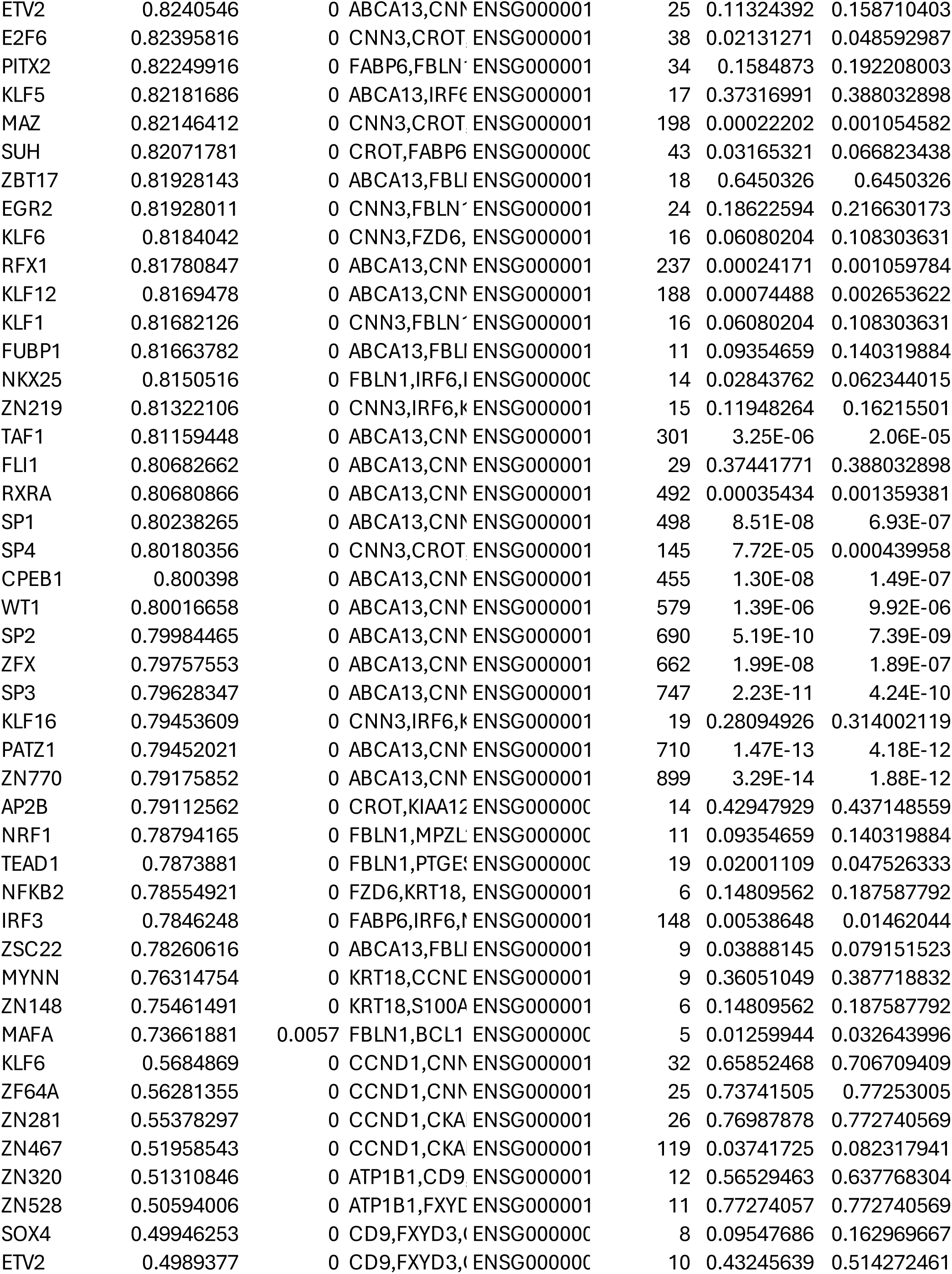
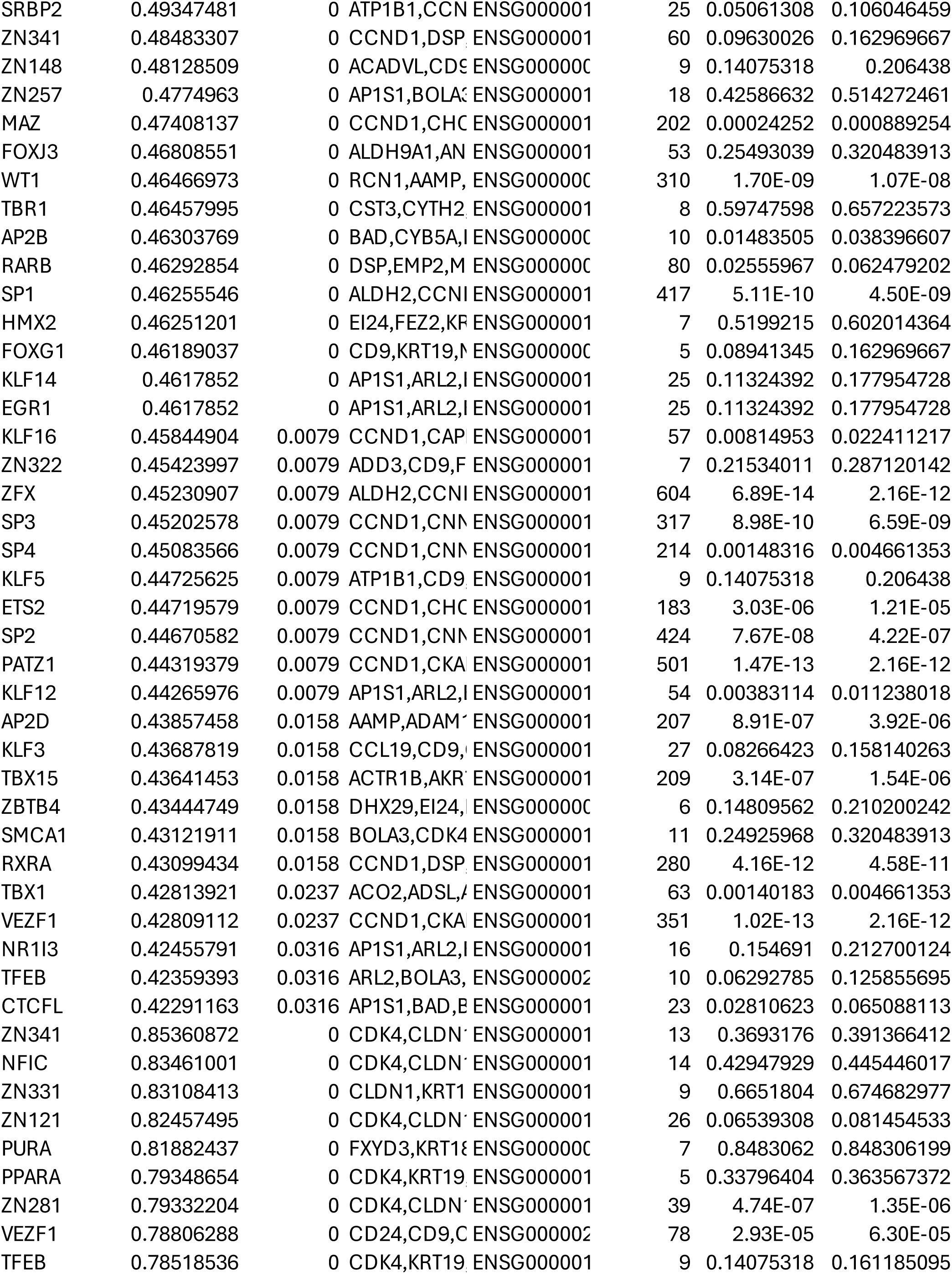
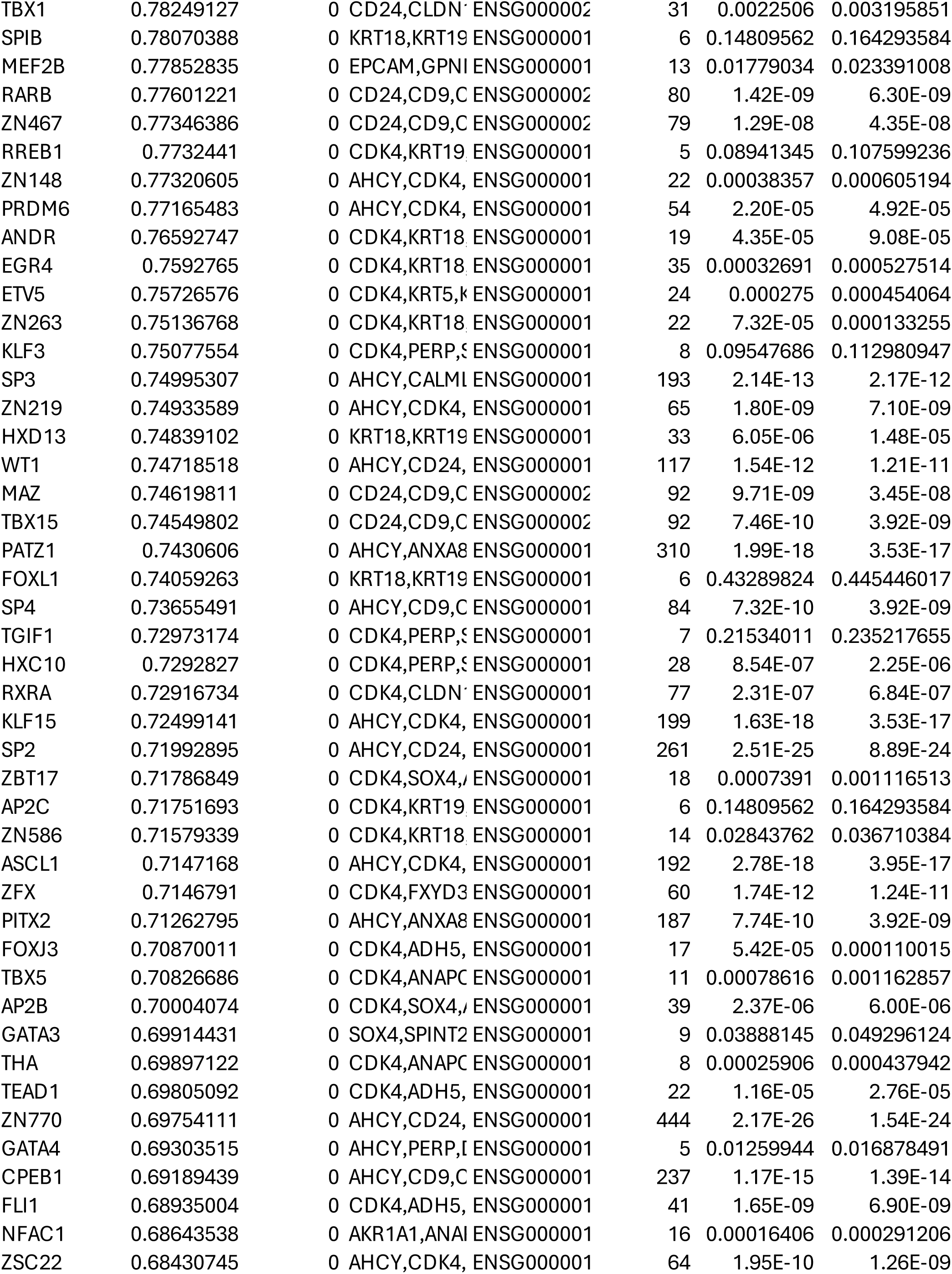
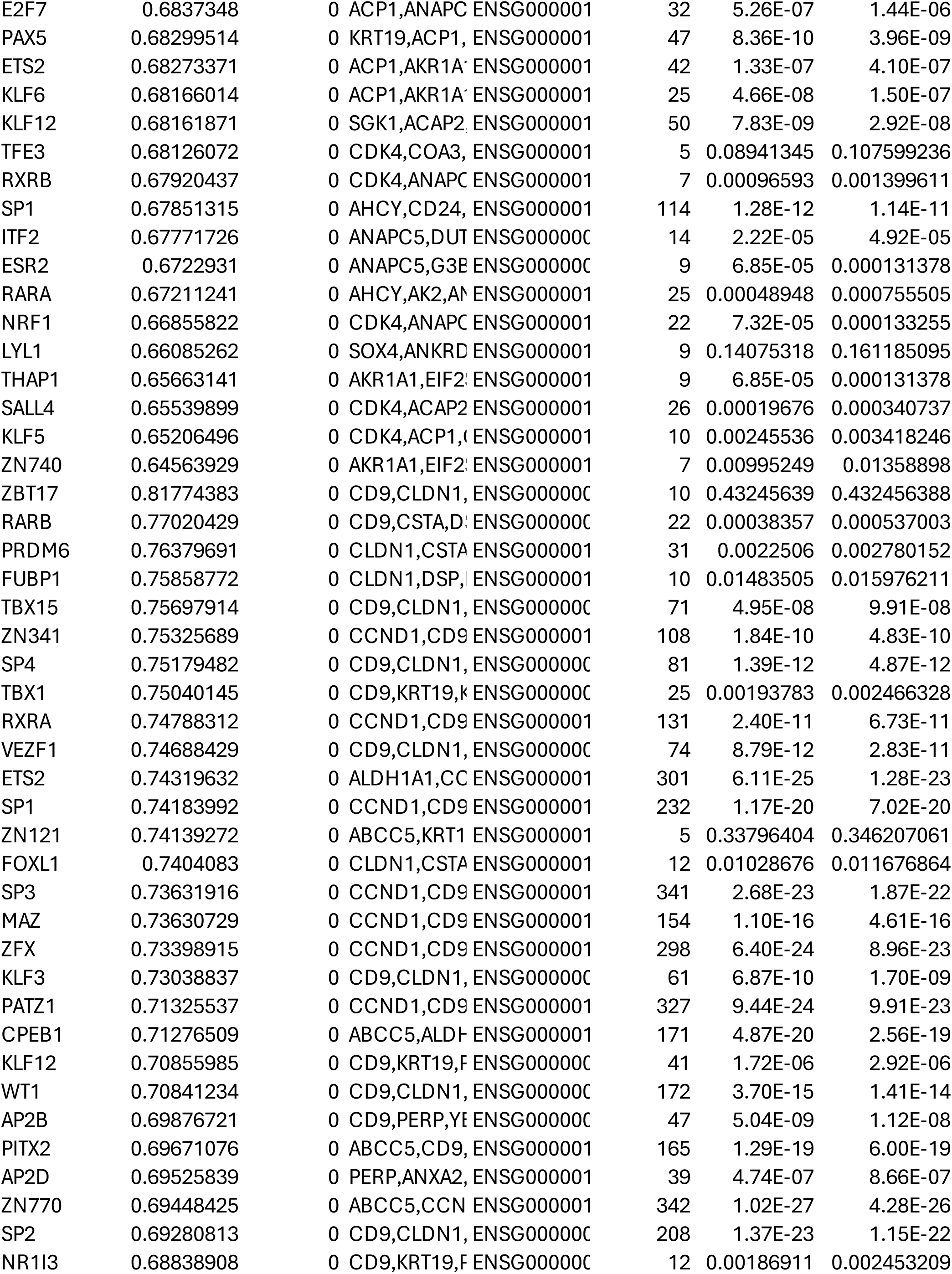
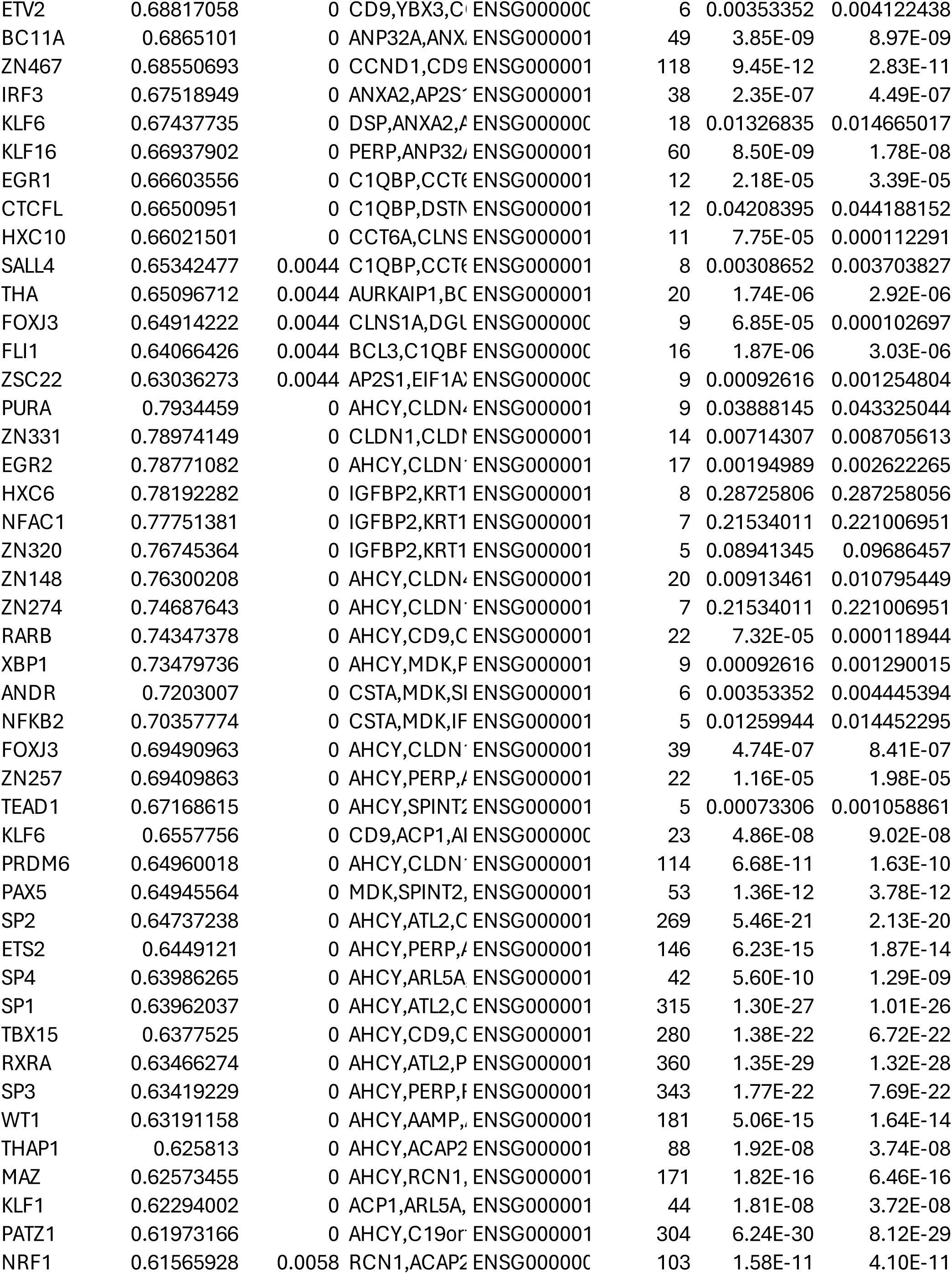
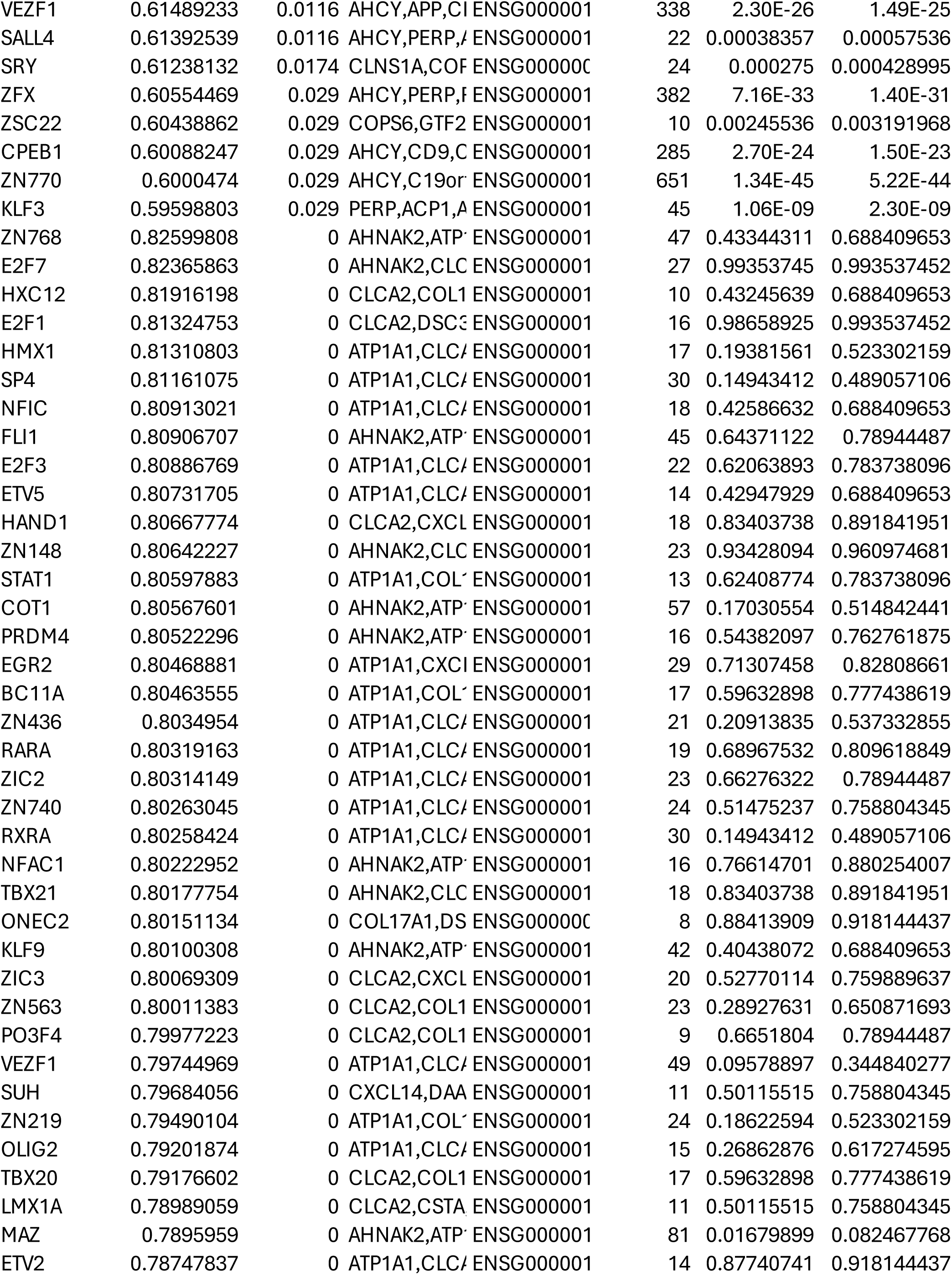
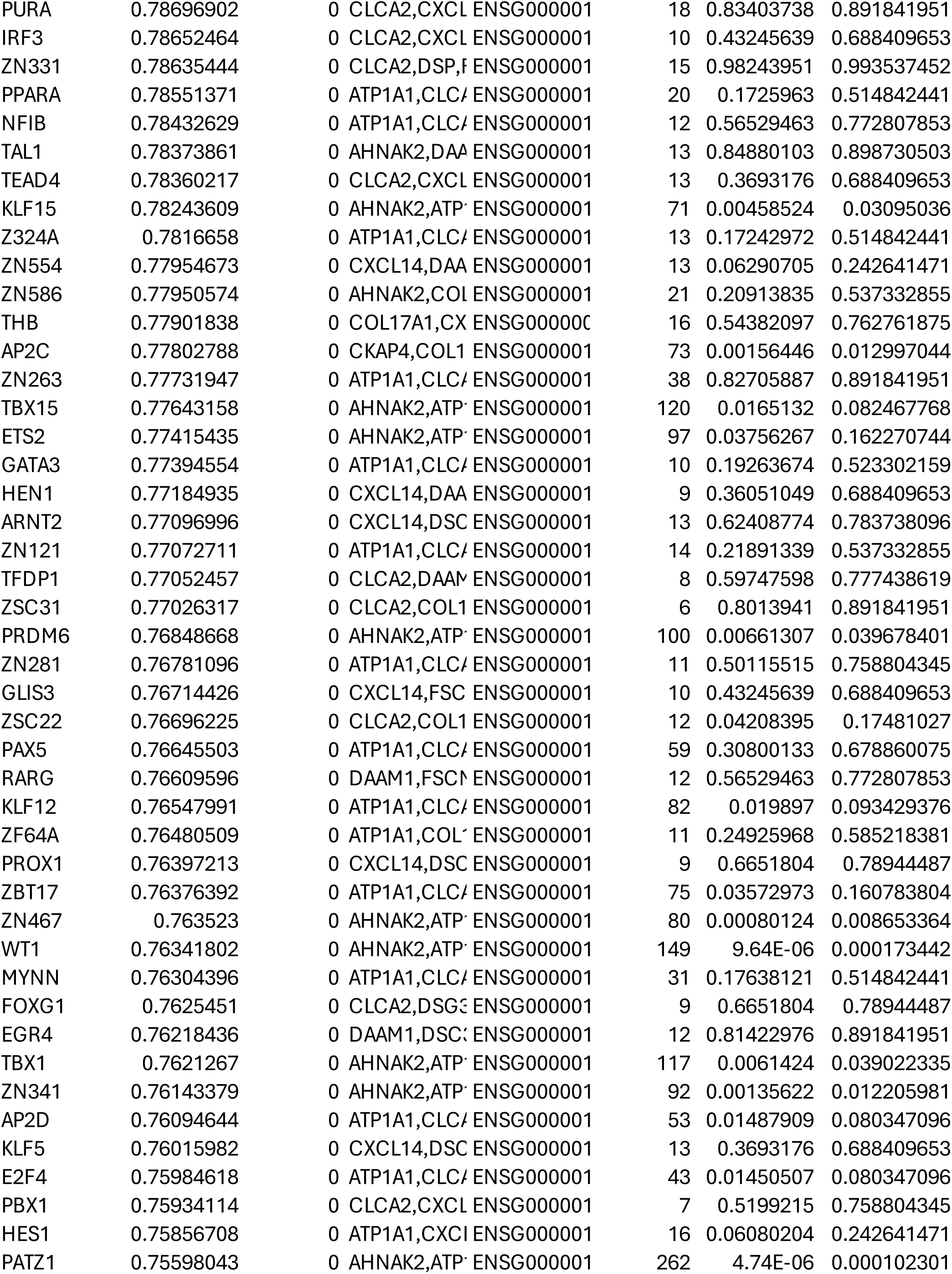
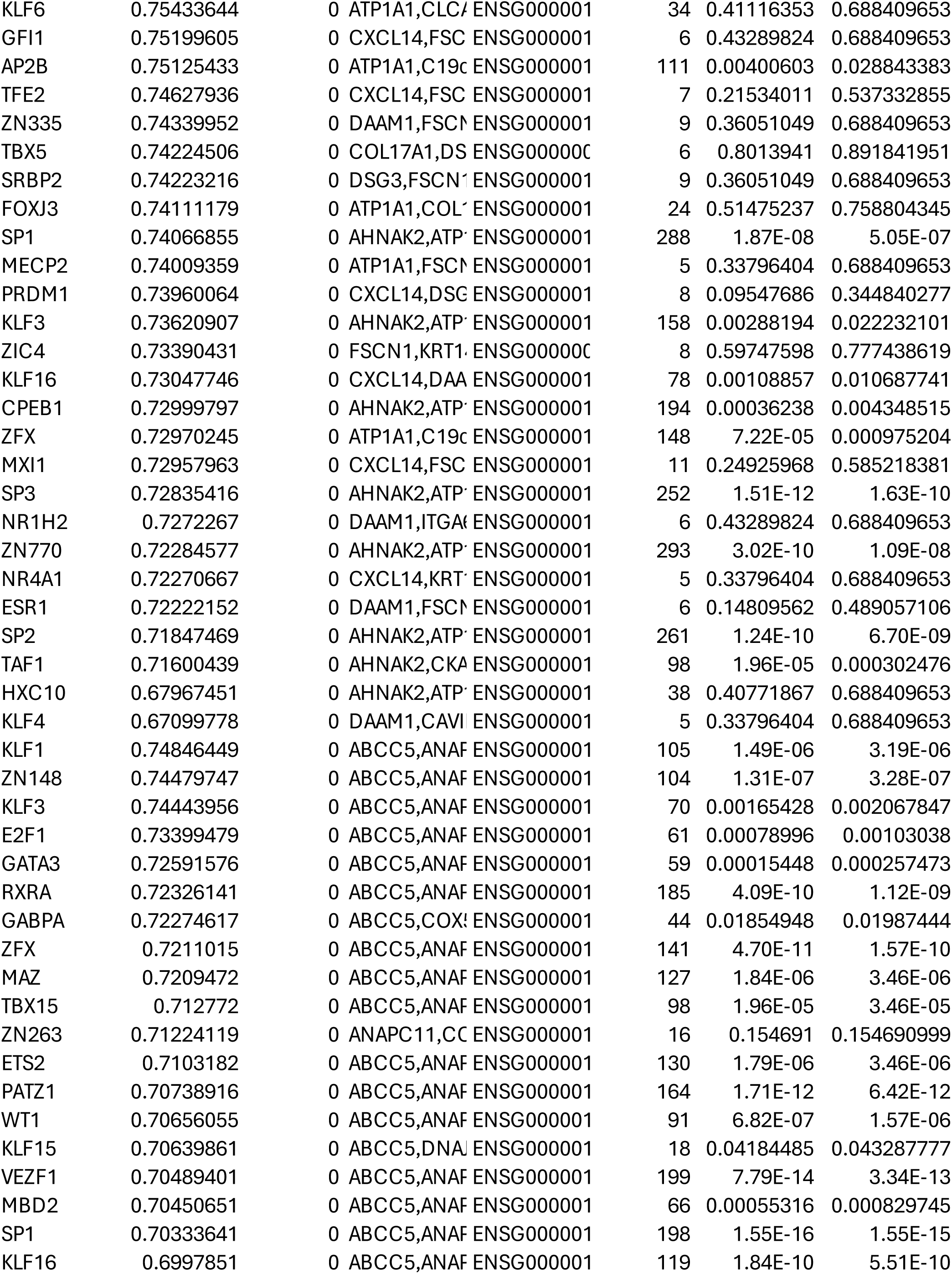
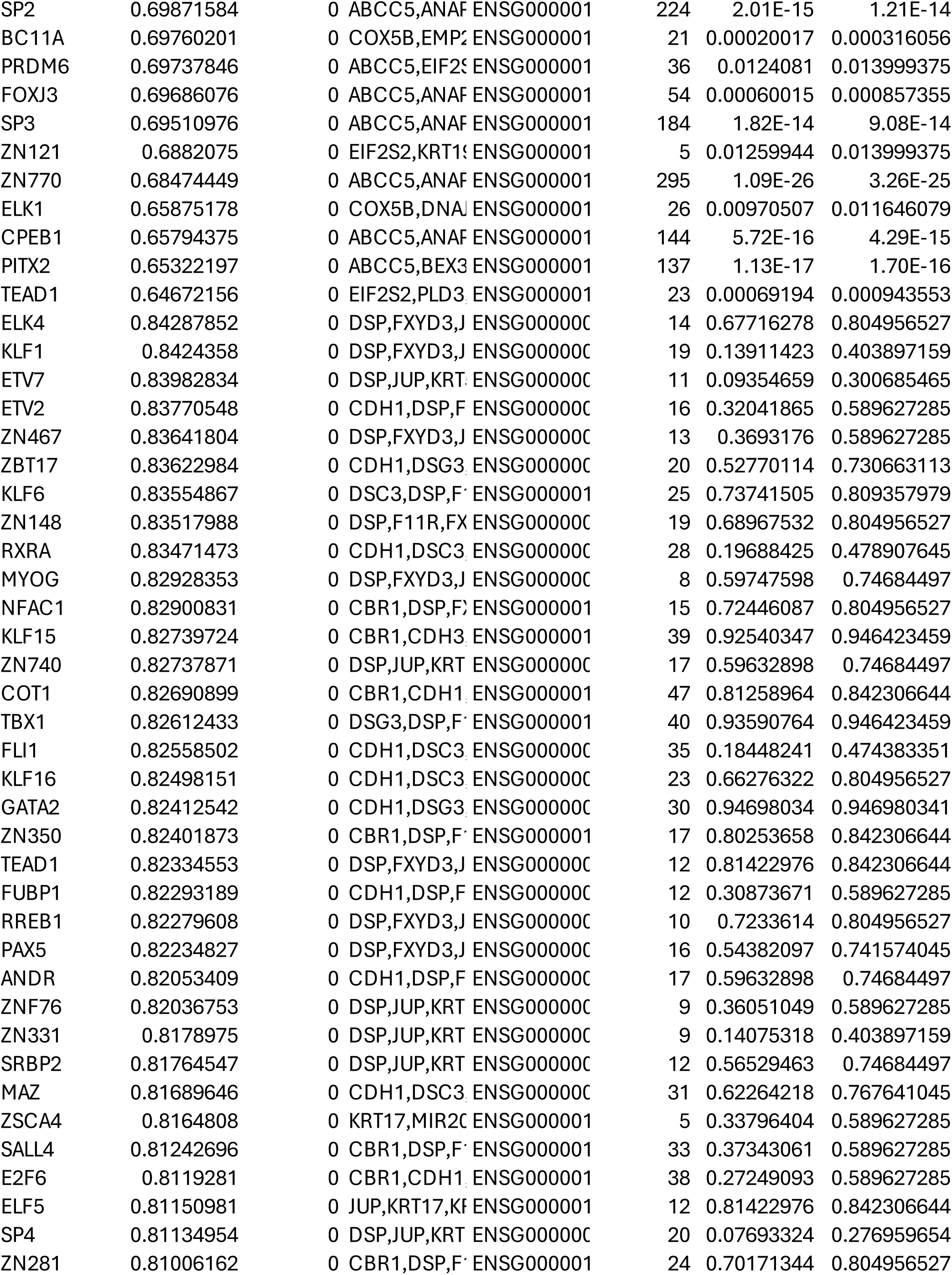
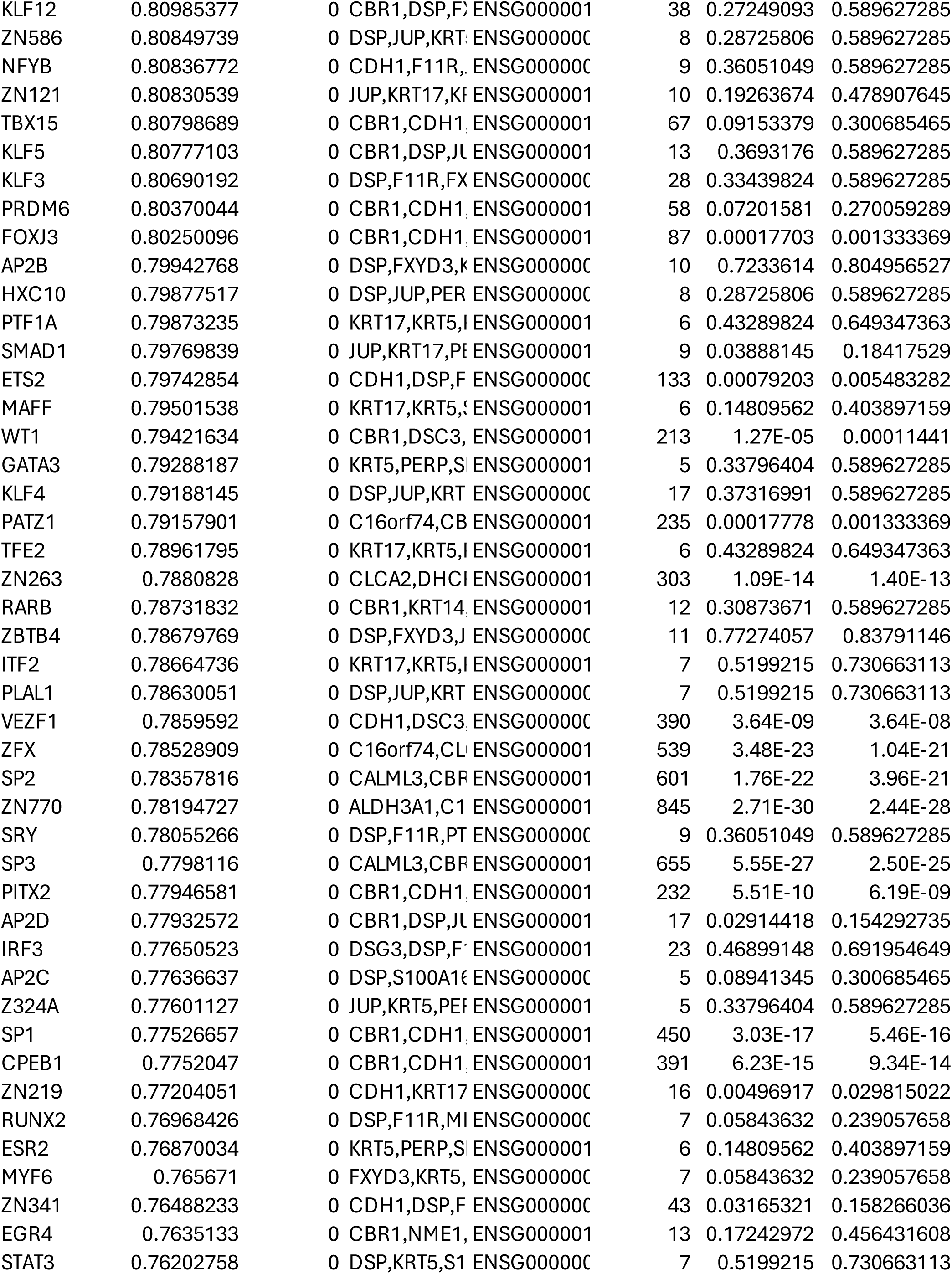
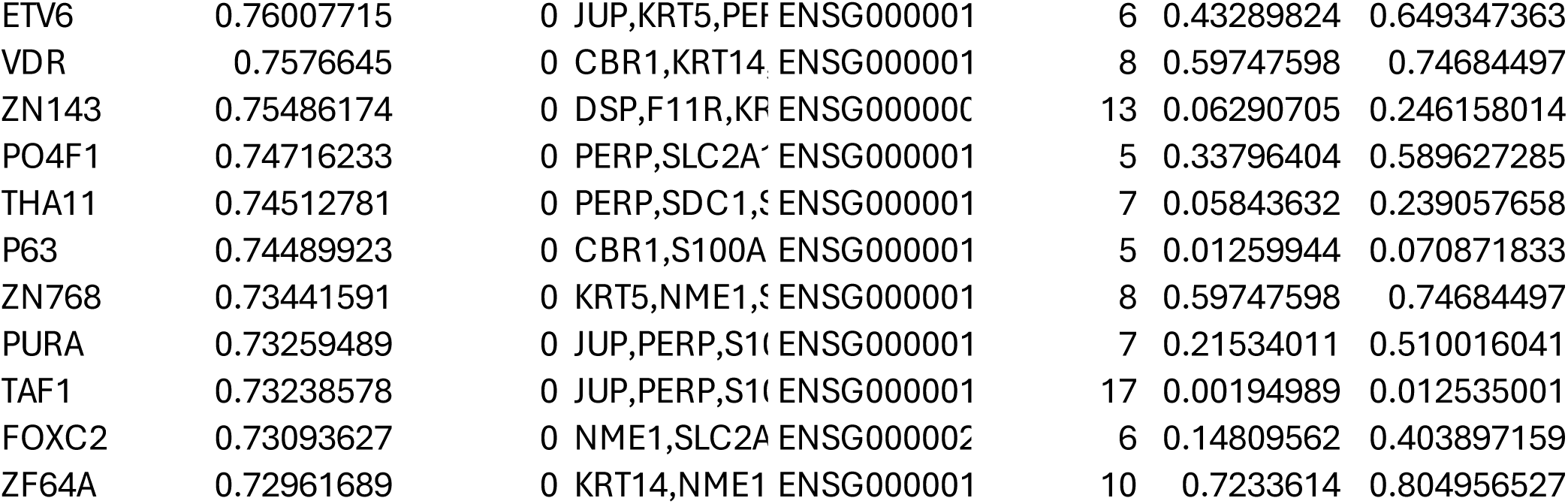
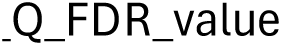
List of 639 TFs that regulate gene expression in HNSCC tumor cells.

**Table S2.**
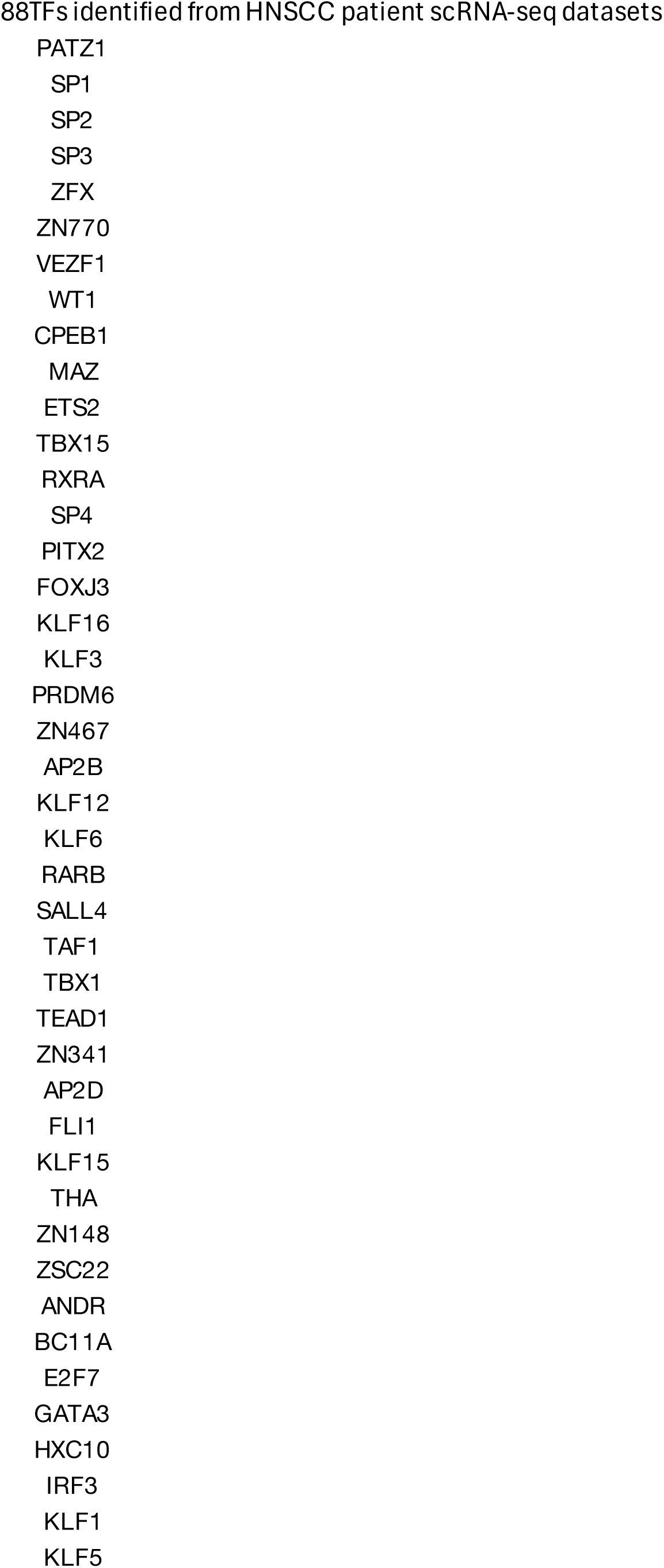
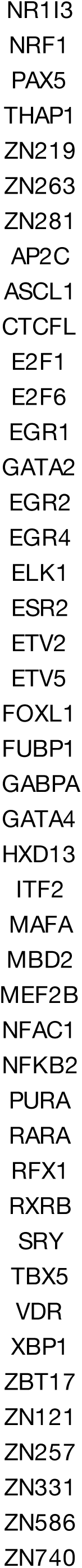
List of 88 TFs that target IRG expression in HNSCC tumor cells identified from at least one patient.

**Table S3.**
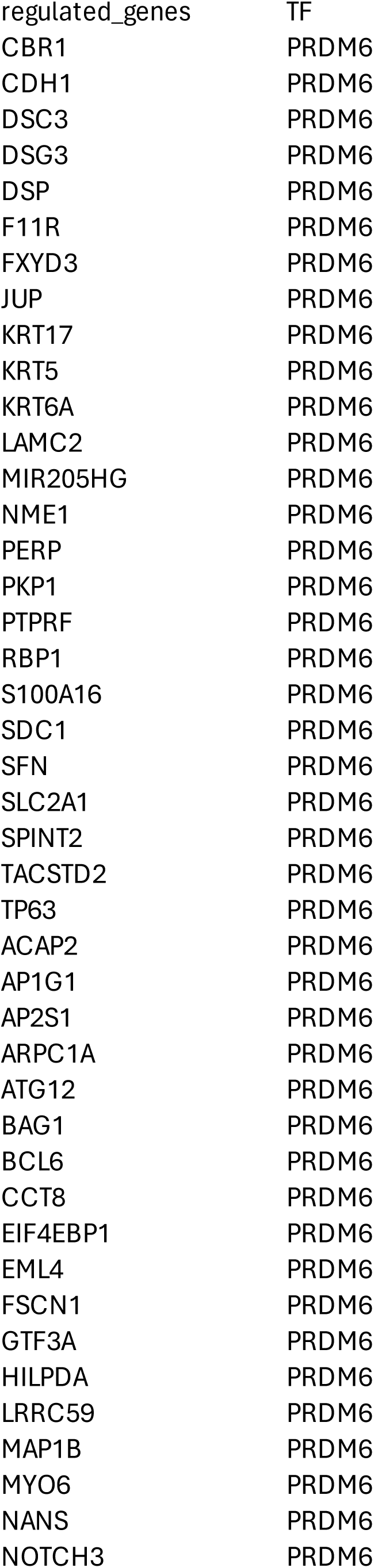
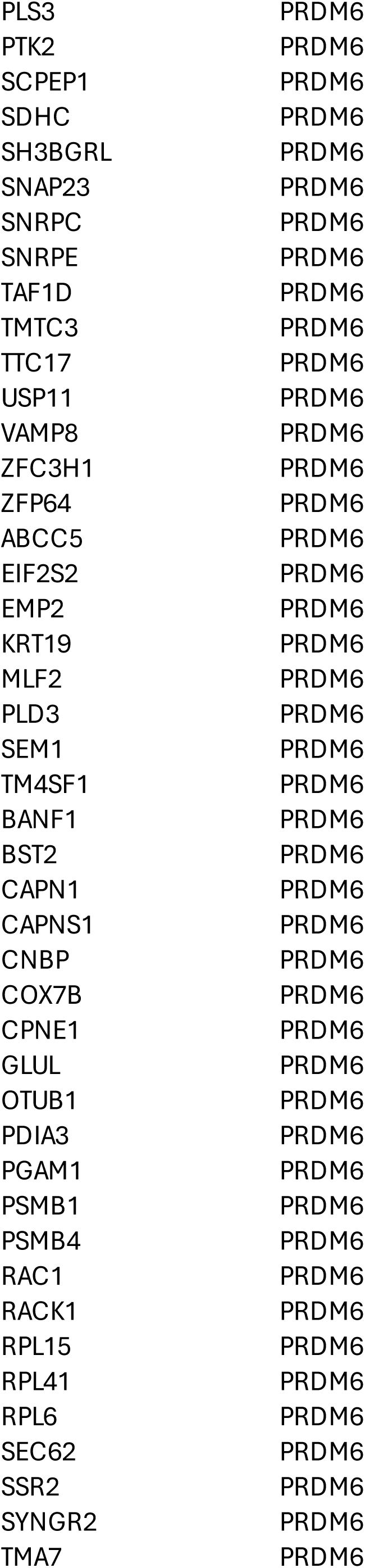
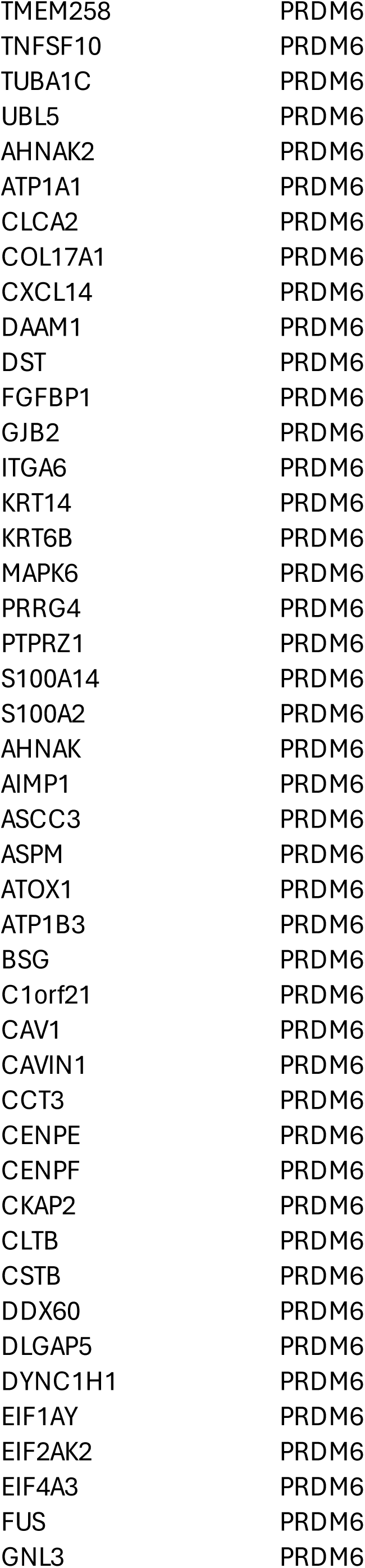
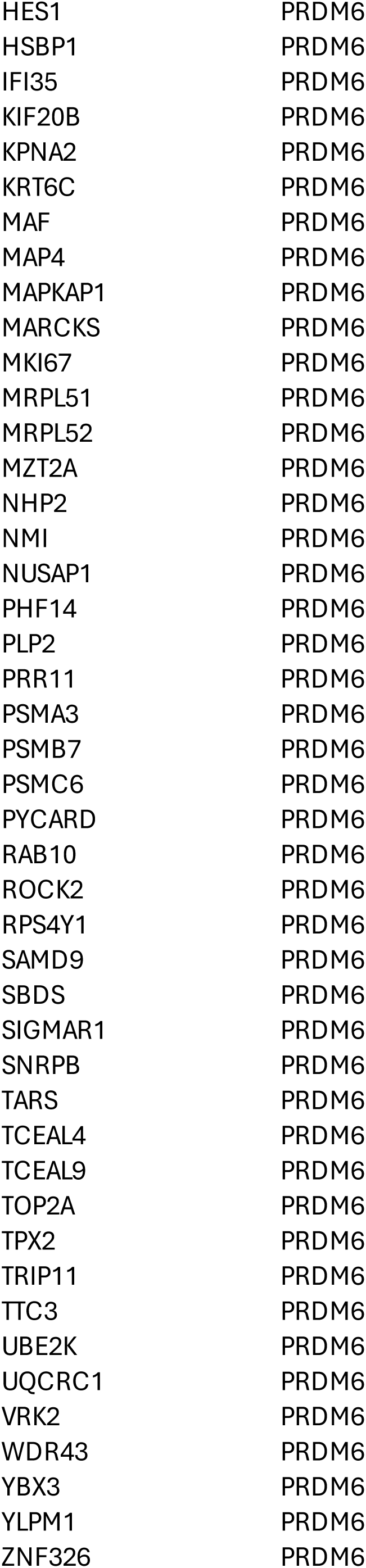
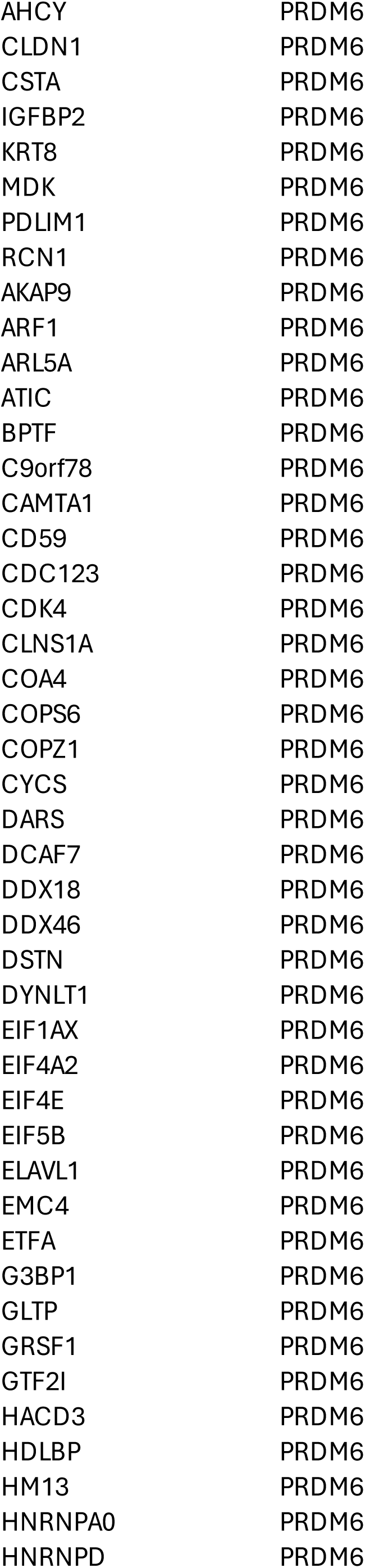
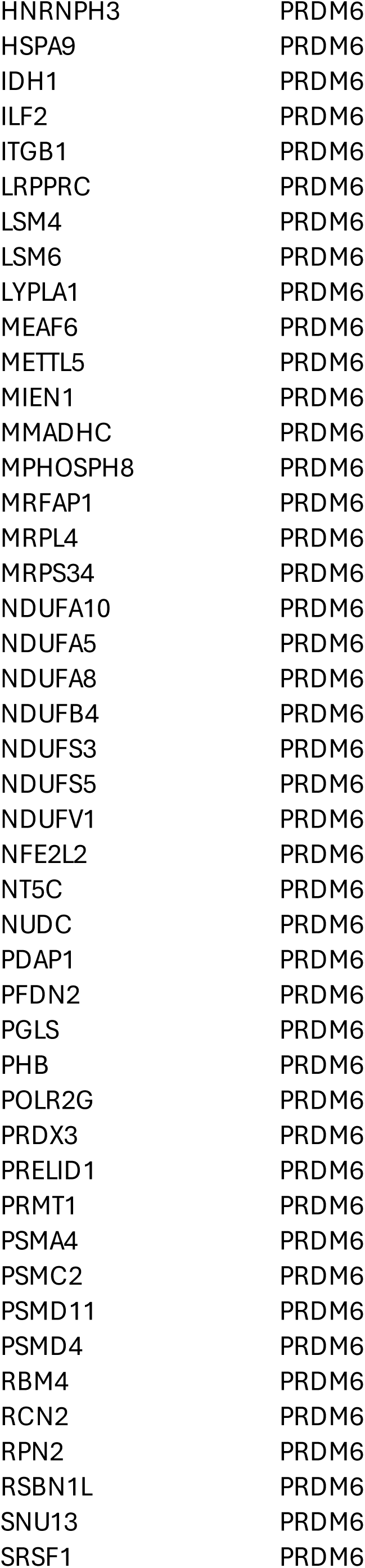
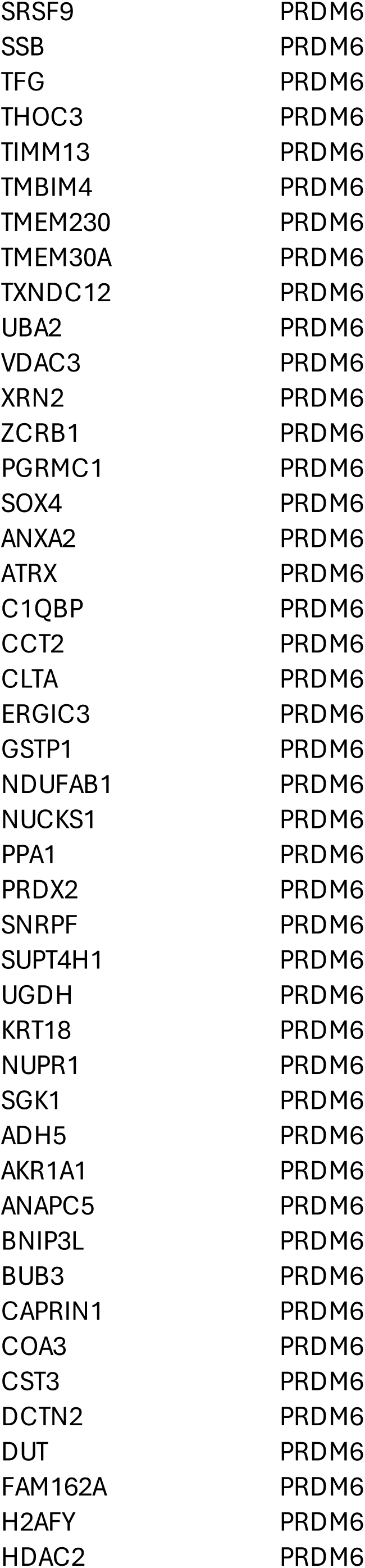
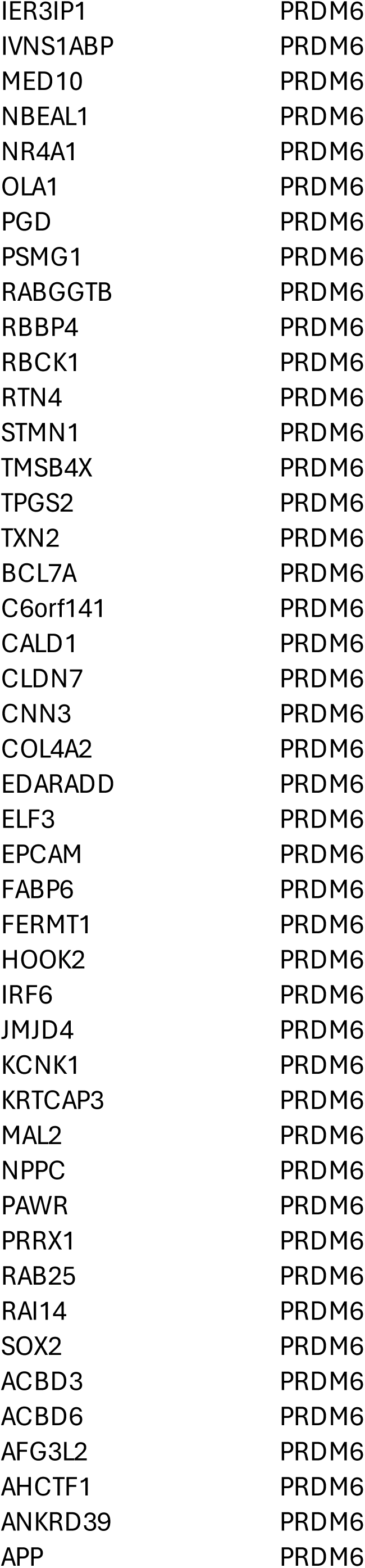
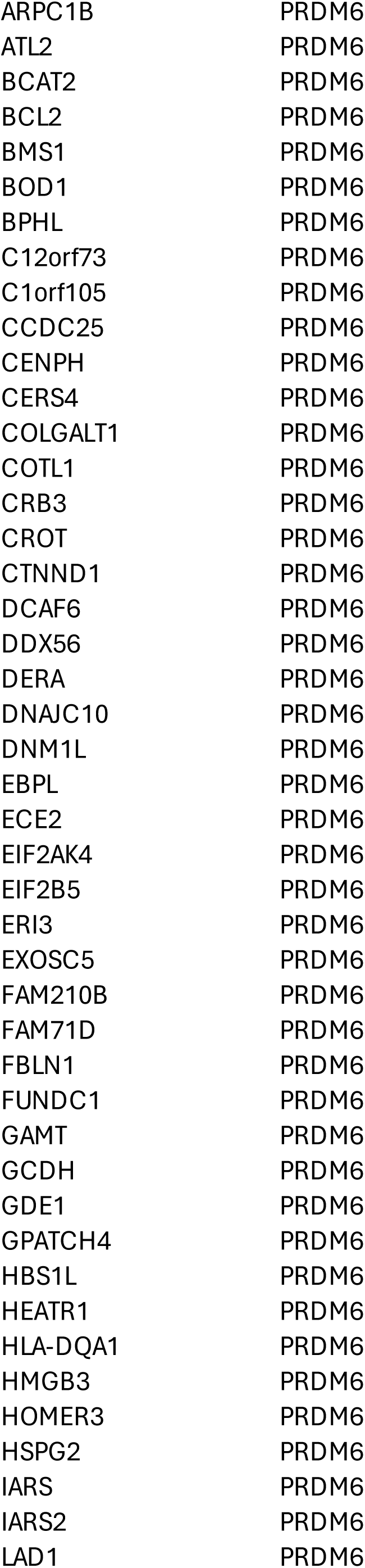
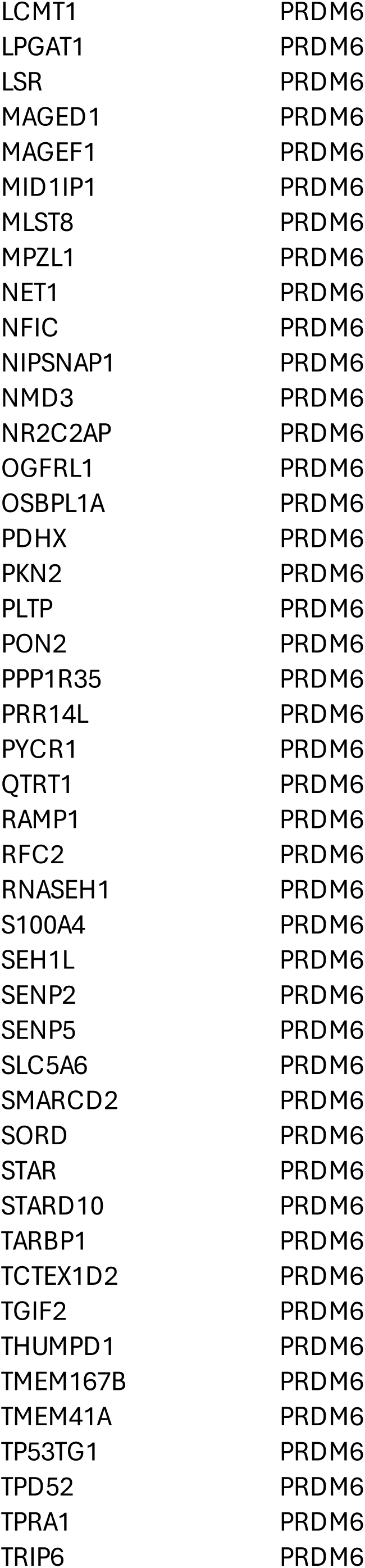
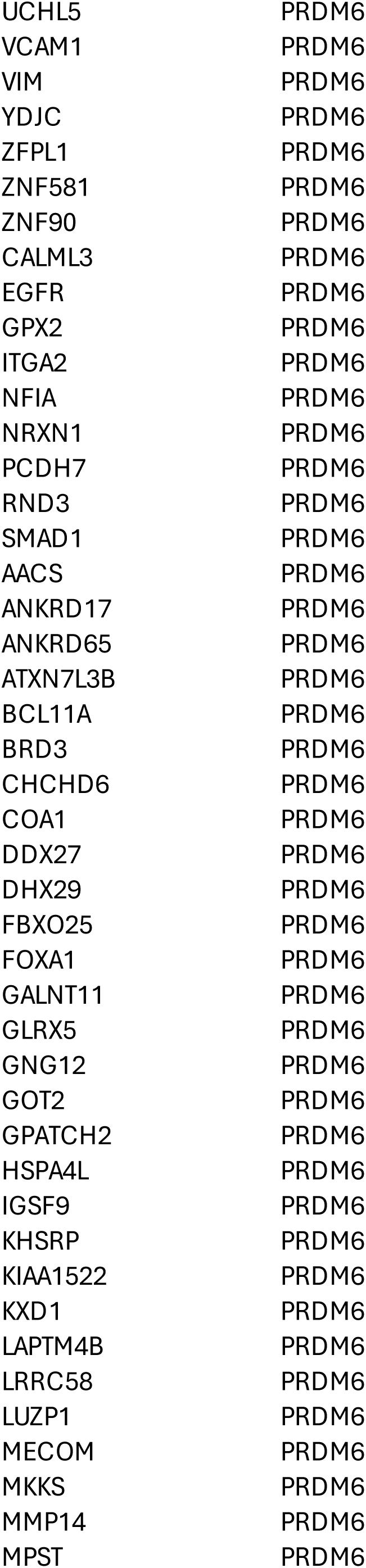
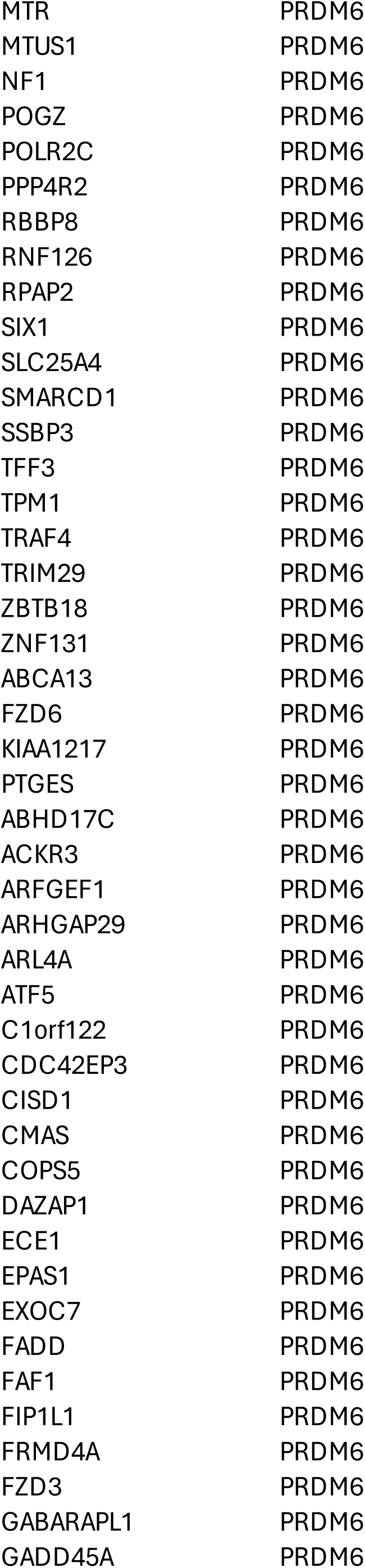
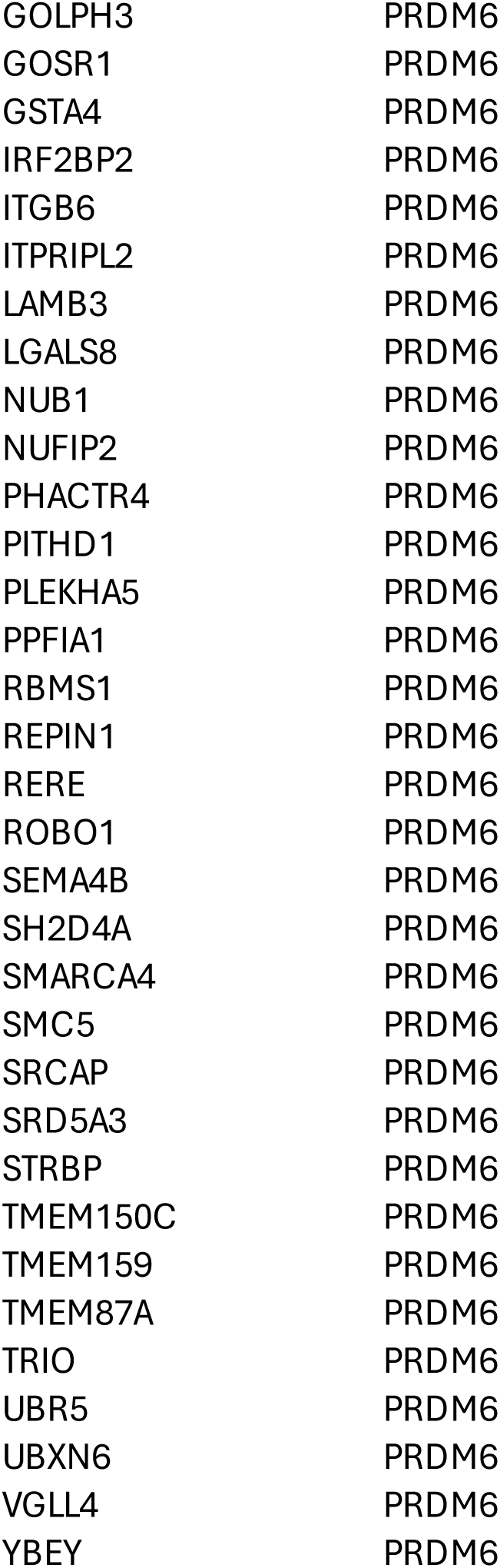
List of 571 genes under regulation of PRDM6.

**Table S4.**
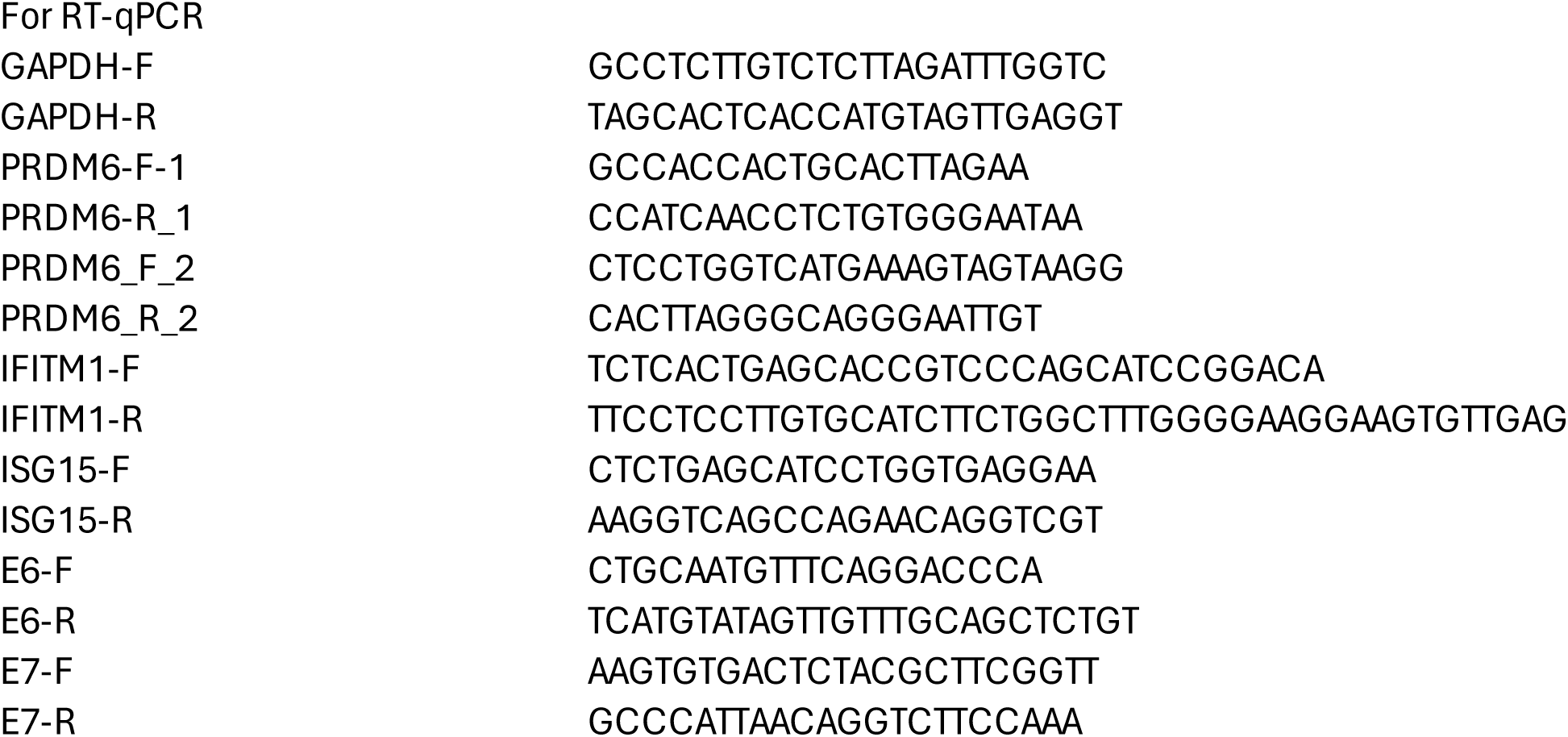
qPCR primers used in the study.

